# Replication-associated solo-WCGW hypomethylation reflects cumulative immune activation across diseases

**DOI:** 10.1101/2025.04.25.650620

**Authors:** Mihoko Shimada, Yosuke Omae, Yuki Hitomi, Yoshiko Honda, Tohru Kodama, Makoto Honda, Katsushi Tokunaga, Taku Miyagawa

## Abstract

Global DNA hypomethylation is a hallmark of immune-mediated diseases, yet its regulatory significance remains unclear. Replication-associated loss of DNA methylation at solo-WCGW (W = A/T) CpGs has recently been proposed as a consequence of cell division. Here we systematically investigated genome-wide hypomethylation patterns across seven immune-mediated diseases. Most diseases exhibited global hypomethylation, particularly at solo-WCGW CpGs in transcriptionally repressed regions, potentially reflecting increased immune cell proliferation. By contrast, CpG sites whose methylation levels were associated with cytokine exposure or SNP genotypes were predominantly located in transcriptionally active regions. To investigate whether immunological events driving immune cell proliferation may also be imprinted in transcriptionally active regions, we searched for differentially methylated regions (DMRs) correlated with an index reflecting the extent of solo-WCGW hypomethylation. In narcolepsy, we identified a DMR within the T-cell receptor alpha (TRA) locus, and greater hypomethylation was associated with increased clonality of both TRA and TRB repertoires, with a similar pattern in an independent cohort. In multiple sclerosis, a DMR was also detected within the IGH locus encoding the B-cell receptor. Together, these findings suggest that the hypomethylation index captures the impact of disease-specific immune dynamics, while reflecting a shared epigenetic signature of immune cell proliferation across diseases.

## Introduction

DNA methylation is a key epigenetic mechanism that affects gene expression and genome stability. Several studies analyzing genome-wide methylation patterns have reported global hypomethylation in immune-mediated diseases, particularly autoimmune diseases such as systemic lupus erythematosus (SLE) ^1–6^, rheumatoid arthritis (RA) ^7–10^, systemic sclerosis ^11,12^, and Sjögren syndrome ^13–15^. Previous studies primarily focused on hypomethylations in regulatory regions and predominantly found methylation changes in these regions, such as those of interferon (IFN)-regulated genes and genes associated with the JAK-STAT pathway ^16,17^; however, they do not account for all methylation sites hypomethylated in the disease.

Global hypomethylation has also been reported in various types of cancer ^18,19^. These hypomethylations are primarily found in megabase-scale regions known as partially methylated domains (PMDs), which are characterized by low gene density, low GC content, late replication timing, and localization to repressed chromatin regions associated with the nuclear lamina ^19,20^. Notably, hypomethylation within PMDs shows strong sequence-context dependence. CpG sites lacking neighboring CpGs (“solo” CpGs), particularly in the WCGW sequence context (W = A/T), exhibit pronounced loss of methylation with increasing cell division ^19,20^. In contrast, CpG sites with neighboring CpGs (“social” CpGs) show little or no such decline ^19,21^. Outside PMDs, referred to as highly methylated domains (HMDs), methylation levels also show a modest decrease at solo-WCGW sites with increasing cell division ^21^. These results suggest that some hypomethylation, especially in the solo-WCGW context, observed in cancers arises from passive demethylation due to failures in the replication and maintenance of methylation processes during cancer cell division ^19,21^. However, few data are available regarding whether such passive DNA demethylation occurs in diseases other than cancer. It has also been reported that PMDs are only present in non-tumor cells of proliferative tissues such as peripheral white blood cells and the placenta ^22^, and another study suggested that PMDs arise only in response to environmental stimuli such as carcinogens ^23^.

In this study, we initially found that most of the methylation sites associated with narcolepsy type 1 (NT1) were hypomethylated in the T cells of NT1 patients, and these hypomethylated CpGs were significantly enriched in the solo-WCGW context. Although NT1 is clinically characterized as a hypersomnia, accumulating evidence suggests that immune mechanisms contribute to the loss of orexin neurons and disease onset ^24^. We then explored whether a similar phenomenon occurs in other immune-mediated diseases. We found significant hypomethylation in the solo-WCGW context in all of the immune-mediated diseases examined. We therefore expanded our investigation to elucidating the underlying causes of the observed CpG hypomethylation.

## Results

### Hypomethylation in immune-mediated diseases and its association with solo-WCGW

In our previous study of narcolepsy type 1 (NT1), we reported that disease-associated CpG sites were predominantly hypomethylated in CD4⁺ and CD8⁺ T cells ^25^. To determine whether hypomethylated CpGs accumulate at CpGs that undergo passive demethylation during cell division, we investigated whether these hypomethylated CpGs are particularly enriched in the solo-WCGW sequence context and found that in both T cell types, NT1-associated hypomethylated CpGs were significantly enriched in the solo-WCGW context (CD4^+^: *P* = 2.57E-179, CD8^+^: *P* = 3.13E-21, Fig 1).

**Fig. 1.**
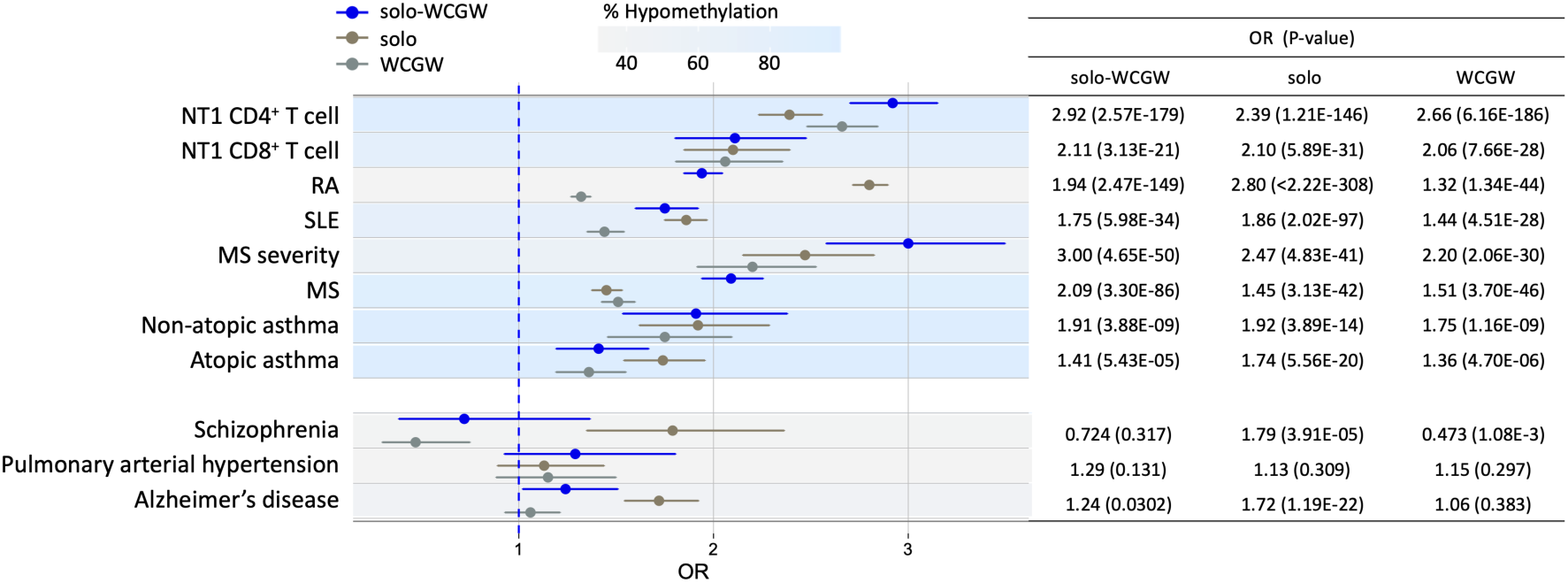
Hypomethylation and the accumulation of hypomethylated CpG sites in the solo-WCGW context in immune-mediated diseases. The proportion of hypomethylated disease-associated CpG sites was examined in immune-mediated and non–immune-mediated diseases. Enrichment of hypomethylated CpG sites in the solo context (i.e., lacking nearby CpGs) and in the WCGW context (CpGs flanked by A or T) was evaluated. In the forest plots, the background color for each disease represents the proportion of hypomethylated CpG sites among all disease-associated CpGs, with darker blue indicating a higher degree of hypomethylation.

To examine whether a similar pattern occurs across other immune-mediated diseases, we analyzed publicly available EWAS datasets including at least 100 cases and 100 controls per study. These datasets included studies of SLE ^26^, RA ^27^, multiple sclerosis (MS) ^28^, MS severity ^29^, atopic asthma ^30^, and non-atopic asthma ^30^ (Table S1). As a comparison, we also analyzed EWAS datasets of non-immune-mediated diseases, including schizophrenia ^31^, pulmonary arterial hypertension ^32^, and Alzheimer’s disease ^33^ (Table S1).

We first examined whether disease-associated differentially methylated positions (DMPs) showed preferential hypomethylation. Marked global hypomethylation was observed in immune-mediated diseases: more than 95.0% of DMPs were hypomethylated in MS, atopic asthma, and non-atopic asthma. Similarly, 74.2% and 57.0% of DMPs were hypomethylated in SLE and MS severity, respectively, whereas only 32.2% were hypomethylated in RA (Fig 1, Table S2). In contrast, hypermethylated sites were more prevalent in all cases of the three non-immune-mediated diseases examined, and no global hypomethylation was observed (Fig 1, Table S2).

We then explored whether these hypomethylations were particularly abundant in the solo-WCGW context. The proportion of solo-WCGW in hypomethylated CpGs was significantly higher in all of the immune-mediated diseases examined (Fig 1, Table S2). This trend was particularly strong in autoimmune diseases and NT1 (*P*<1E-20). In non-immune-mediated diseases, however, only Alzheimer’s disease showed a nominally significant increase in solo-WCGW, but no strong associations similar to those observed in immune-mediated diseases were found (Fig 1, Table S2).

### PMDs in non-cancerous cells and their association with solo-WCGW

Although hypomethylation of solo-WCGW is particularly prominent in PMDs in cell models (Fig 2a), PMDs are reported to be largely absent in non-dividing cells ^23,34^. We therefore examined PMDs in immune cell subsets using an established method ^19,21^. We detected PMDs in 15 non-cancerous immune cell subsets, as well as in neurons and tumor samples for comparison (Table S3) ^19,35^. The tumor samples exhibited particularly large regions detected as PMDs (Fig S1a) and PMDs exhibited cell/tissue specificity (Fig S1b, S1c). Further, PMDs in tumors were approximately 10 times longer than in non-cancerous cells (Fig S1d and S2). Tumor samples harbored 5–10 times more PMD regions than immune cells, with only 32.7% overlapping those in immune cells (Table S4). Furthermore, PMDs in tumor samples were predominantly found in transcriptionally repressed regions with heterochromatin and outside of CpG island-associated regions, whereas in non-cancerous cells, PMDs were located more frequently in regions with open chromatin, transcriptionally active regions, and CpG islands (Fig 2b and S3). These results suggest that in non-cancerous cells, local methylation variability at transcriptional regulatory regions may have been identified as PMDs, indicating that PMDs with typical features such as low gene density and late replication timing, as seen frequently in cancer, are largely absent in non-cancerous cells. Therefore, as expected, when we examined whether the hypomethylated solo-WCGW CpGs observed in immune-mediated diseases were enriched in the PMDs of immune cells, except for non-atopic asthma, the odds ratios (ORs) for solo-WCGW sites in immune-cell PMDs were not higher than those for overall solo-WCGW sites, and no significant enrichment was found (Fig 2c, Table S5).

**Fig. 2.**
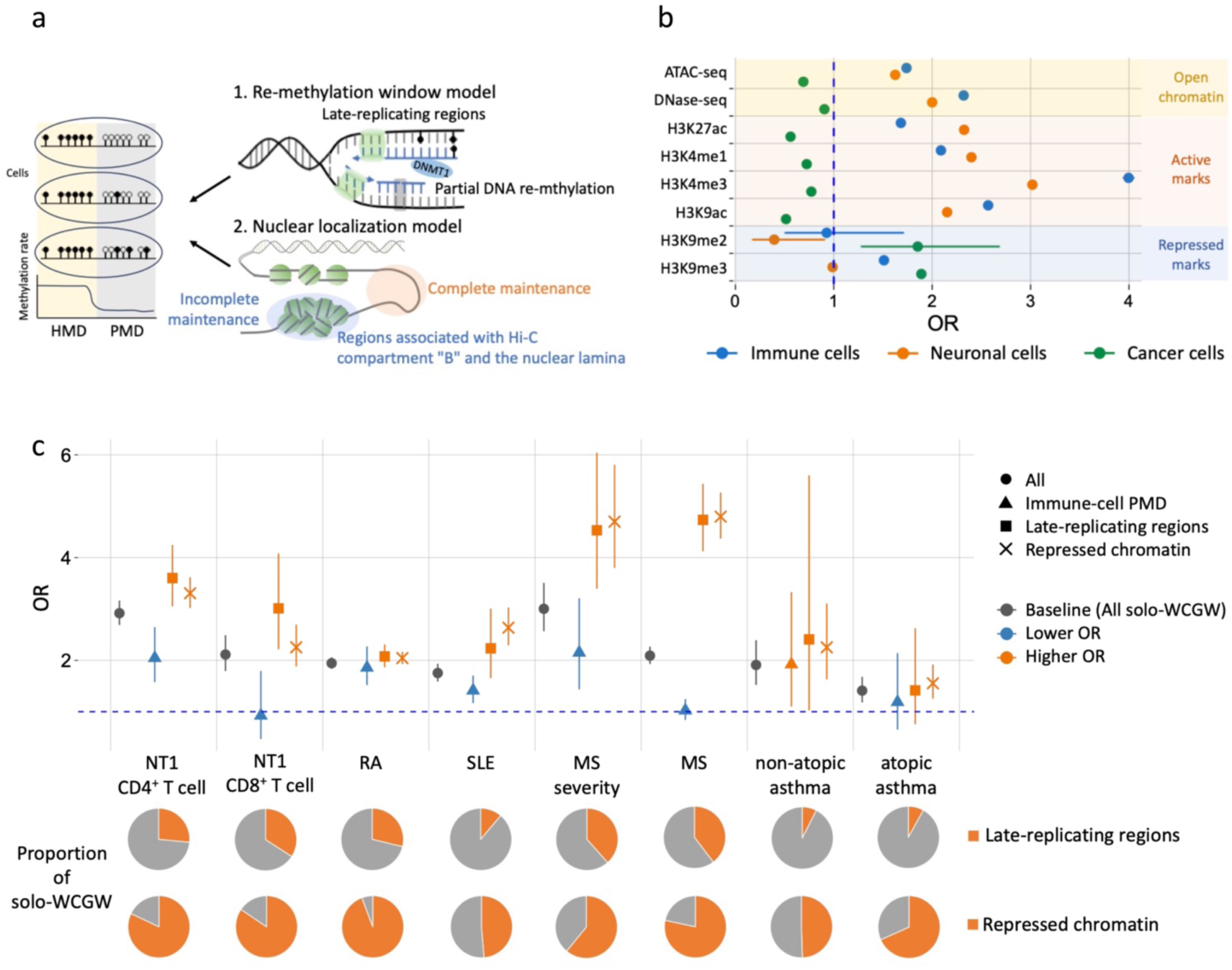
Identification and characterization of regions in which passive demethylation of solo-WCGW is likely to occur. **a.** Schematic models proposed for passive demethylation of solo-WCGW sites. **b.** Overlap between the regions identified by ATAC-seq, DNase-seq, and histone ChIP-seq in immune cells and the PMDs in cancer, neural, and immune cells. **c.** Comparison of the odds ratios (ORs) of the presence of disease-associated solo-WCGW in immune-cell PMDs, regions with late replication timing, and repressed chromatin. For comparison of the ORs of all disease-associated solo-WCGW in Fig 1, the ORs of disease-associated solo-WCGW in each region are depicted in orange when higher and blue when lower. Pie charts show the proportion of disease-associated solo-WCGW present in late-replicating regions and repressed chromatin.

### Solo-WCGW sites in immune-mediated diseases are enriched in late-replicating domains and repressed chromatin

In previous *in vitro* cell model studies, PMDs were found predominantly in late-replicating domains, in which passive demethylation is thought to occur during replication, a concept known as the re-methylation window model (Fig 2a) ^19^. We therefore next examined the relationship between replication timing and solo-WCGW hypomethylation observed in immune-mediated diseases. Using publicly available Repli-seq data for non-cancerous B cells, we examined whether hypomethylated solo-WCGW sites are enriched in late-replicating regions. In all immune-mediated diseases, the ORs for hypomethylated solo-WCGW in late-replicating regions were higher than those of hypomethylated solo-WCGW overall (Fig 2c, Table S6). However, hypomethylated solo-WCGW in late-replicating regions accounted for only 7.5–39.7% of all hypomethylated solo-WCGW in each disease (Fig 2c).

We next examined the relationship between hypomethylated solo-WCGW sites and closed chromatin regions such as lamina-associated domains, proposed as another explanation for passive methylation loss during cell division (the nuclear localization model; Fig 2a) ^19^. To assess the distribution of hypomethylated solo-WCGW sites across these chromatin features, we classified CpG sites as transcriptionally active euchromatin or other non-active regions (repressed chromatin) using integrated ATAC-seq, DNase-seq, and histone modification data. Not only were the ORs for solo-WCGW sites in repressed chromatin higher than those for hypomethylated solo-WCGW overall, but these sites also accounted for a larger proportion of hypomethylated solo-WCGW CpGs than regions with late replication timing (25.3–86.0%, Fig 2c, Table S7). These results suggest that although late-replicating regions are important, passive demethylation associated with replication is particularly likely to occur in repressed chromatin, especially in immune-mediated diseases.

Regarding NT1, we previously obtained methylation data from whole blood-derived DNA using a 27K array ^36^. Using these data, we determined whether the same result observed in the analysis of CD4^+^ and CD8^+^ T cells was reproducible. In whole blood, a trend toward hypomethylation was also observed at methylation sites associated with NT1 (Table S8). Hypomethylated sites were significantly enriched in repressed chromatin (OR = 2.81, *P* = 2.44E-06) and more frequently found in solo CpGs lacking surrounding CpGs (OR = 2.28, *P* = 5.51E-04); however, they were not particularly enriched in solo-WCGW (OR = 0.976, *P* = 1.00) (Table S8).

### Effect of cytokines on changes in global methylation

Hypomethylation in the context of solo-WCGW is not necessarily caused solely by passive demethylation along with increased cell proliferation; for instance, in immune-mediated diseases, active demethylation may occur due to the effects of various cytokines ^37^. Therefore, we explored whether TNF-α and TGF-β, two cytokines implicated in immune diseases, induce hypomethylation, and whether they affect the hypomethylation of solo-WCGW, particularly in repressed chromatin. We utilized data from previous studies in which cells were treated with the cytokines, and changes in methylation sites after treatment were compared with pre-treatment ^38,39^. First, we examined the direction of methylation changes in CpG sites affected by the cytokines. With TNF-α treatment, the majority of CpG sites (86.5%) shifted toward hypomethylation (Fig 3a), whereas with TGF-β treatment, the majority of CpG sites (90.3%) shifted toward hypermethylation (Fig 3b). We then examined whether hypomethylated CpGs were particularly enriched in solo-WCGW within repressed chromatin. With TNF-α treatment, although solo-WCGW sites were significantly enriched (OR = 2.00, *P* = 8.00E-26, Table S9), hypomethylated CpGs were predominantly found in open chromatin regions with transcriptionally active histone marks (DNase-seq: OR = 7.33, *P*<2.22E-308, H3K27ac: OR = 2.48, *P* = 9.59E-65) (Fig 3a, Table S10). Although hypomethylated CpGs represented a small fraction of all CpGs, those associated with TGF-β treatment were found in open and transcriptionally active chromatin regions in cells treated with TGF-β (ATAC-seq: OR = 6.61, *P*<2.22E-308, H3K27ac: OR = 4.93, *P*<2.22E-308, Table S10), whereas hypomethylated CpGs were less common in the solo-WCGW context (OR = 0.105, *P* = 1.04E-76) (Fig 3b, Table S9). These results suggest that at least as pertaining to the two cytokines examined in this study, hypomethylated CpGs induced by these cytokines occur predominantly in open chromatin regions and do not sufficiently explain the solo-WCGW hypomethylation in repressed chromatin observed in immune diseases.

**Fig. 3.**
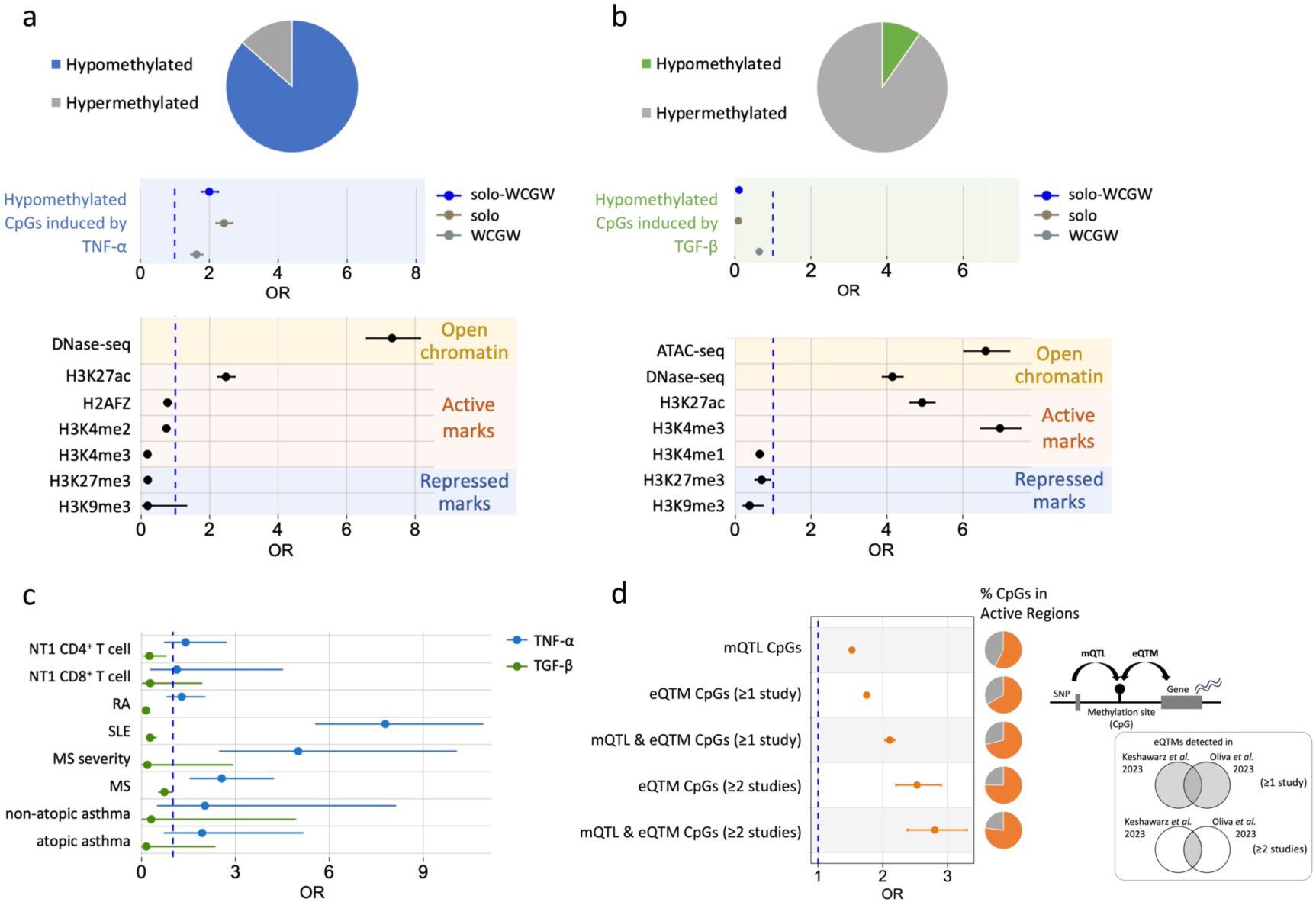
Hypomethylation associated with cytokines and genetic variants is enriched in transcriptionally active regions. **a.** Changes in DNA methylation upon TNF-α treatment in human umbilical vein endothelial cells. The pie chart shows the proportion of CpG sites undergoing hypomethylation among those with altered methylation after TNF-α treatment. The middle forest plot shows the ORs for solo-WCGW among hypomethylated CpG sites. The bottom forest plot shows the ORs for enrichment of hypomethylated CpG sites in genomic regions identified by DNase-seq and histone ChIP-seq analyses in endothelial cells. **b.** Changes in DNA methylation upon TGF-β treatment in ovarian cancer cell lines. Similar analyses were performed as in **a**. Bottom panel shows a forest plot based on regions identified by ATAC-seq, DNase-seq, and histone ChIP-seq in ovaries. **c.** Odds ratios (ORs) for the overlap between disease-associated hypomethylated CpG sites and CpG sites hypomethylated in response to cytokine stimulation. **d.** ORs calculated by comparing the proportions of CpG sites with mQTLs and/or eQTMs located in transcriptionally active regions versus repressed chromatin. Pie charts show the proportion of CpGs located in mQTLs, eQTMs, or both that fall within active regions. For eQTMs, “≥1 study” indicates analyses using eQTMs detected in either of the two studies (upper Venn diagram), whereas “≥2 studies” indicates analyses using eQTMs detected in both studies (lower Venn diagram).

Subsequently, we investigated the overlap between CpGs hypomethylated by TNF-α and TGF-β and disease-associated hypomethylated CpGs. Whereas CpGs hypomethylated by TGF-β did not significantly overlap with disease-associated hypomethylated CpGs in any of the diseases examined, CpGs hypomethylated by TNF-α overlapped significantly with hypomethylated regions in SLE, MS, and MS severity (Fig 3c, Table S11).

### Transcriptionally active-region enrichment of genotype- and expression-associated CpGs

Next, we examined CpGs associated with methylation quantitative trait loci (mQTLs) and expression quantitative trait methylation (eQTMs). These CpGs, whose methylation levels are influenced by genetic variants or correlate with gene expression, were analyzed for their distribution between transcriptionally active and repressed regions as defined in our study. CpGs located in mQTLs identified from large-scale whole-blood samples in a previous study ^40^ were significantly enriched in transcriptionally active regions (OR = 1.52, *P* <2.22E-308, Fig 3d, Table S12). For eQTMs, we used data registered in the eQTM-Catalog derived from two previous studies in which whole blood was analyzed ^41,42^. eQTM CpGs were significantly enriched in active regions, and, in particular, eQTM CpGs detected in both studies exhibited a stronger enrichment in active regions (OR = 2.53, *P* = 1.33E-44, Fig 3d, Table S12). Furthermore, CpGs that were both influenced by SNPs and associated with gene expression—that is, CpGs overlapping mQTLs and eQTMs—showed even greater enrichment in active regions (OR = 2.80, *P* = 5.21E-41, Fig 3d, Table S12).

### Active-region DMRs associated with the global solo-WCGW hypomethylation index

The degree of solo-WCGW hypomethylation showed strong concordance across the genome within individuals (Fig S4), indicating that it can be represented by a global hypomethylation index. We therefore searched for differentially methylated regions (DMRs) in transcriptionally active chromatin whose methylation levels correlate with this hypomethylation index. Analyses were performed in the NT1 and MS datasets, for which genome-wide single-CpG methylation data were available. In the NT1 analyses, the solo-WCGW hypomethylation index correlated with estimated naïve T-cell proportions (Fig S5), consistent with its association with immune cell proliferation. We then searched for DMRs associated with the solo-WCGW hypomethylation index after adjusting for estimated naïve T-cell proportions. No DMRs replicated at FWER<0.05 were detected in the CD8⁺ T-cell analysis (Table S13, S14). In contrast, the DMR in the T cell receptor alpha (TRA) locus showing the strongest association in the CD4⁺ T-cell analysis was replicated in an independent sample set (FWER<0.05) (Table 1, Fig 4a, Table S15).

**Fig. 4.**
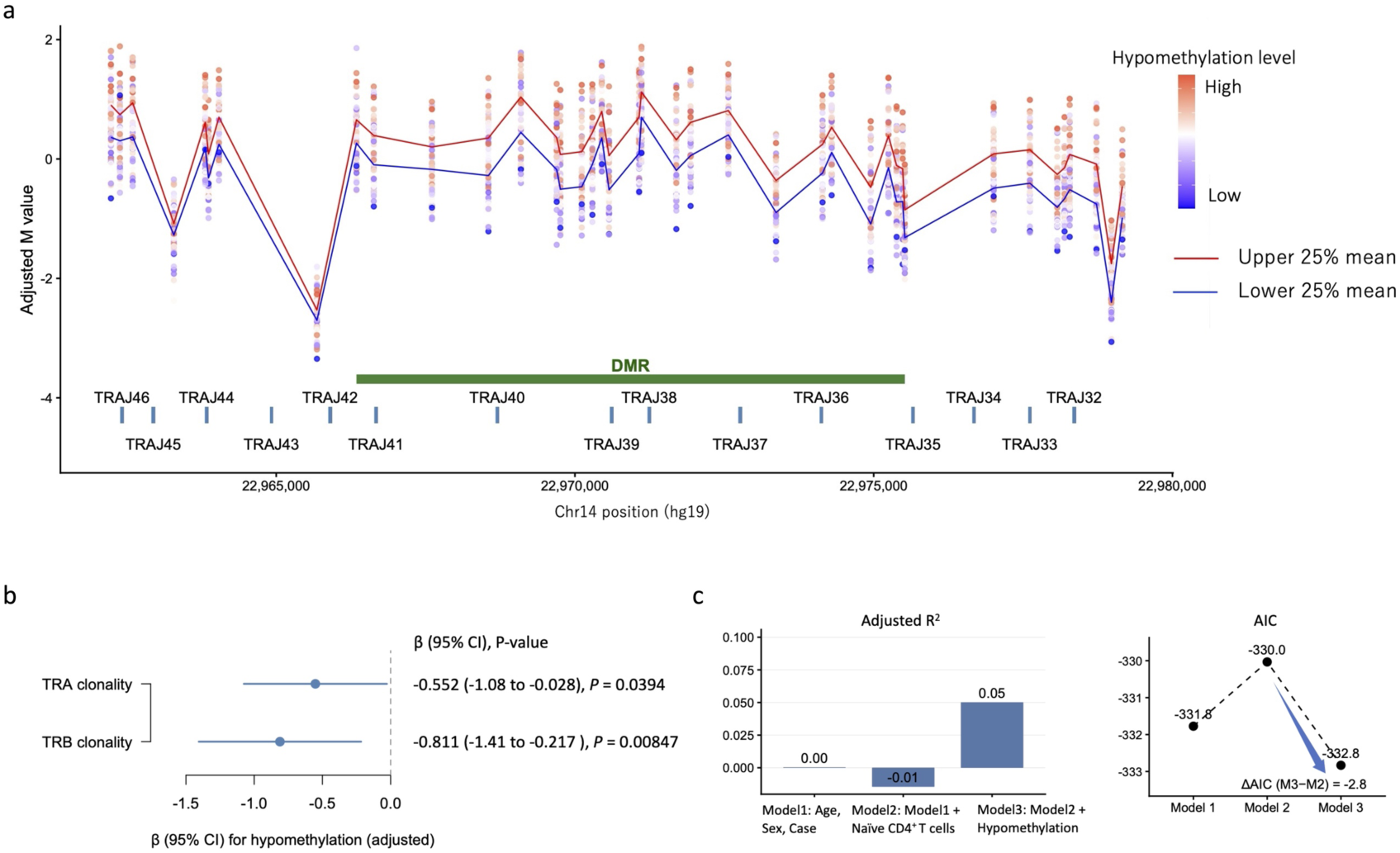
Hypomethylation index correlates with methylation changes at the TRA locus and T-cell receptor clonality in NT1 CD4^+^ T cells. **a.** Methylation levels across the TRAJ locus on chromosome 14 in CD4⁺ T cells from NT1 samples. Points represent adjusted methylation values for individual CpG sites (covariates: age, sex, disease status, naïve CD4⁺ T-cell proportion), colored by hypomethylation level. Samples were stratified by the hypomethylation index, and lines indicate the mean methylation levels for the upper and lower quartiles. The green bar indicates the differentially methylated region (DMR) associated with the hypomethylation index. **b.** Association between the hypomethylation index and T-cell receptor clonality estimated from RNA-seq data. Effect sizes (β) with 95% confidence intervals are shown for TRA and TRB clonality. **c.** Comparison of regression models explaining T-cell clonality. Inclusion of the hypomethylation index improved model fit, as indicated by the increase in adjusted R² and the reduction in AIC.

**Table 1.** DMRs associated with hypomethylation index in NT1 CD4^+^ T cells.

| DMR | Chr | Position (hg19) |  | Value | L | P-value | FWER | Replication |  | Genes in eQTM |
| --- | --- | --- | --- | --- | --- | --- | --- | --- | --- | --- |
|  |  | start | end |  |  |  |  | FWER<0.05 | P<0.05 |  |
| DMR1 | chr14 | 22966344 | 22975521 | 34.8 | 24 | 0 | 0 | TRUE | TRUE | TRDV2:TRDC:TRAJ37:TRAV1-2 |
| DMR2 | chr15 | 98503878 | 98504173 | -71.3 | 4 | 1.09E-05 | 0.068 |  | TRUE |  |
| DMR3 | chr17 | 46656572 | 46660097 | -33.9 | 22 | 2.26E-05 | 0.15 |  | TRUE | HOXB2:HOXB3:HOXB4:HOXB-AS1 |
| DMR4 | chr6 | 42928079 | 42928920 | -31.2 | 20 | 2.56E-05 | 0.17 |  | TRUE | GNMT:PEX6:AL035587.1 |
| DMR5 | chr20 | 36148604 | 36149022 | -28.5 | 20 | 2.59E-05 | 0.172 |  | TRUE |  |

**Abbreviations:** DMR, differentially methylated region; FWER, family-wise error rate; eQTM, expression quantitative trait methylation; L, number of CpGs within the DMR.

We next estimated T-cell receptor clonality using RNA-seq data derived from the same samples. TCR repertoires reconstructed with TRUST4 ^43^ revealed that samples with higher levels of solo-WCGW hypomethylation exhibited increased clonality in both the TRA (*P* = 0.0394) and TRB repertoires (*P* = 0.00847) (Fig 4b). To assess whether the hypomethylation index explains variation in T-cell clonality beyond known covariates, we compared regression models with sequential covariate adjustment. Incorporating the hypomethylation index increased the adjusted R² from -0.01 to 0.05 and resulted in the lowest AIC among the models tested (ΔAIC = -2.8), indicating improved model fit (Fig 4c).

In the MS analysis, although an independent replication dataset was not available, 15 DMRs were identified at FDR<0.2. Among these, a DMR located within the immunoglobulin heavy-chain (IGH) locus was detected (Table S16).

### Prioritizing disease-associated genes through eQTM mapping

Given that chromatin states may differentially influence regulatory potential and that hypomethylation in repressed chromatin appears to capture a cumulative footprint of immune activation, we next asked whether EWAS signals are preferentially enriched in active regulatory regions. Using eQTM associations, disease-associated CpG sites were linked to gene expression, only when genes were supported by multiple CpGs meeting the specified criteria (Fig S6). As a result, no candidate genes were identified in repressed regions across all diseases, reinforcing the limited functional impact of methylation loss in silent chromatin. In contrast, our analysis of transcriptionally active regions revealed disease-associated genes in SLE and MS (Table S17, Table S18, Fig S7a and S7b). The STRING analysis ^44^ indicated that genes negatively correlated with SLE-associated hypomethylated CpG sites were significantly enriched in the type I interferon (IFN-α/β) signaling pathway (Fig S7a), which had not been reported in the original EWAS of SLE ^26^. Among these protein-coding genes, 80.6% have been implicated in SLE in large-scale whole-blood gene-expression studies ^45,46^ (Table S17). As illustrated by the *IFITM1* example in Fig S7c, multiple CpG sites in the eQTMs correlated with *IFITM1* were associated with SLE, consistent with increased *IFITM1* expression in SLE. Similarly, STRING analysis of genes whose expression is negatively associated with hypomethylated CpG sites in MS demonstrated significant enrichment in interferon signaling pathway (Fig S7b). Some of these genes had been reported to be associated with Type I interferon therapy in MS (Table S18).

## Discussion

In this study, we identified five key characteristics related to hypomethylation in immune-mediated diseases (Fig 5). First, hypomethylation was observed in most immune-mediated diseases, with disease-associated hypomethylation sites being particularly abundant in the solo-WCGW context (Fig 1). Second, such hypomethylated solo-WCGW sites were significantly more abundant in late-replicating regions and repressed chromatin (Fig 2). Third, cytokine-induced hypomethylation occurred primarily in open chromatin (Fig 3), and CpGs in mQTLs and eQTMs were also enriched in transcriptionally active regions (Fig 4a). Fourth, the hypomethylation index of solo-WCGW sites in repressed chromatin was associated with the *TRA* locus in NT1 and correlated with T-cell clonality. In MS, the index was associated with the *BCR* locus, collectively suggesting that it captures an epigenetic footprint of cumulative immune activation. Finally, integrating disease-associated hypomethylated CpGs with eQTM information highlighted disease-associated genes and pathways when analyses were restricted to transcriptionally active regions, including the type I interferon signaling pathway in SLE (Fig S7). These results suggest that hypomethylation in repressed chromatin largely reflects indirect effects of immune activation through cell proliferation, whereas hypomethylation in active regulatory regions more directly captures disease-associated regulatory changes (Fig 5).

**Fig. 5.**
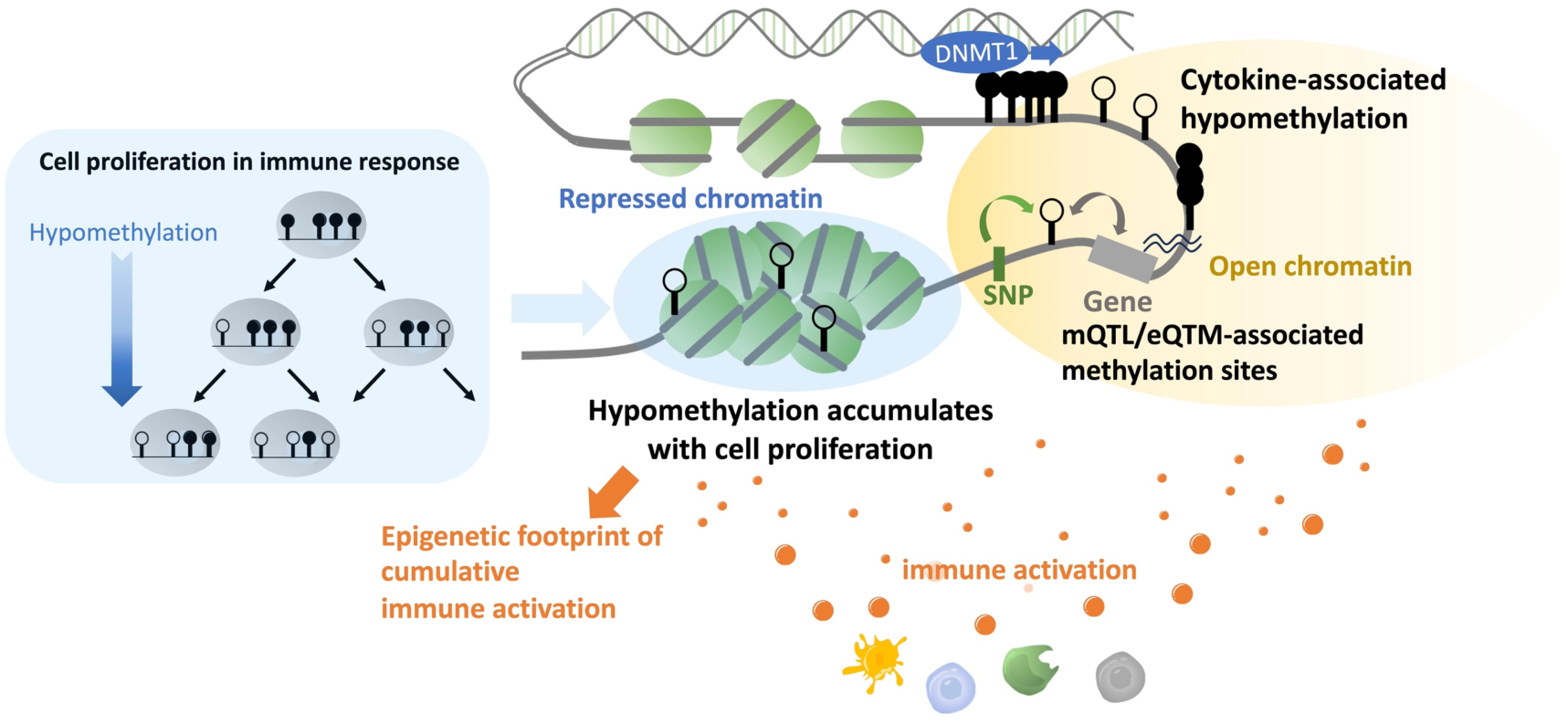
Model of replication-associated hypomethylation as an epigenetic footprint of cumulative immune activation. Immune stimulation directly influences methylation in transcriptionally active regions, where regulatory changes such as cytokine signaling and genetic effects occur. In contrast, CpG sites in repressed chromatin are less directly affected by regulatory processes. Instead, immune activation promotes immune cell proliferation, leading to replication-associated passive demethylation that accumulates particularly at solo-WCGW sites. Because repressed chromatin is less exposed to other regulatory influences, this hypomethylation can accumulate as an epigenetic footprint of cumulative immune activation.

Previous studies have shown that passive demethylation occurs at CpG sites within the solo-WCGW sequence context following cell division ^19,21^. In this study, we observed a consistent trend across all immune-mediated diseases examined. Just as cell proliferation is enhanced in cancer, the proliferation of immune cells, including T cells and B cells, is promoted during immune responses ^47,48^. Other studies have shown that eosinophilia in asthma involves not only mobilization from the blood but also proliferation and differentiation of progenitor cells in the lungs ^49,50^. Thus, the hypomethylation of solo-WCGW observed in immune-mediated diseases can be attributed to the enhanced proliferation of immune cells involved in the pathology. Several mechanisms have been proposed to explain the increased susceptibility to CpG demethylation in the context of solo-WCGW. One hypothesis suggests that because DNMT1 (the enzyme responsible for maintaining DNA methylation patterns) is a processive enzyme, once it binds to a CpG it efficiently methylates surrounding CpGs ^51,52^. Another hypothesis suggests that the presence of nearby methylated CpGs increases the likelihood that repair mechanisms will address demethylation ^53,54^. However, the affinity of DNMT1 for CpGs in the WCGW context remains controversial ^55,56^, and further research is needed to clarify this issue.

Previous studies have shown that passive demethylation of solo-WCGW sites associated with cell division occurs primarily within PMDs, especially prominent in cancerous cells ^19,21^; however, this was not the case in the immune-mediated diseases examined in the present study. This discrepancy is likely attributable to fundamental differences in the characteristics of PMDs between cancer cells and non-malignant immune cells. In cancer cells, large-scale hypomethylated regions are clearly detected as PMDs. In contrast, in immune cells, transcriptional regulatory regions can be detected as PMDs, suggesting that normal immune cells may not harbor PMDs as prominent as those observed in cancer cells. In immune-mediated diseases, hypomethylated solo-WCGW sites are predominantly found in regions with late replication timing, supporting the re-methylation window model. However, because the total number of analyzed CpGs located in late-replicating regions is relatively small, the overall proportion of solo-WCGW sites in these regions remains limited. Instead, a greater proportion of solo-WCGW sites can be attributed to repressed chromatin (Fig 2c). These results suggest that the nuclear localization model may also be valid, particularly in the case of immune-mediated diseases.

Previous studies have reported global hypomethylation in autoimmune diseases such as SLE, with the accumulation of hypomethylated regions in IFN-related genes ^16,17^. Indeed, it is possible that immune factors such as cytokines contribute to the occurrence of hypomethylation. However, in the present study, CpGs exhibiting hypomethylation following treatment with the cytokines TNF-α or TGF-β were located primarily in open chromatin. This is consistent with previous studies showing that upon B cell stimulation, the majority of methylation changes occur in the direction of demethylation, predominantly affecting accessible chromatin and enhancer regions ^57,58^. Given the limited sample size for TGF-β (n = 3) and the small number of cytokines examined, further investigation is warranted. Nevertheless, our findings suggest that cytokine effects alone are insufficient to explain hypomethylation in repressed chromatin in immune-mediated diseases. Similar to the hypomethylation induced by cytokines, our analysis revealed that CpGs in mQTLs and eQTMs are enriched in transcriptionally active regions, indicating that focusing on transcriptionally active regions may not only enhance the interpretation of DNA methylation and gene expression analyses, but also facilitate the identification of genetic variants associated with traits in genome-wide association studies. Indeed, when we used eQTM relationships to search for genes whose expression is suggested to be linked to disease-associated hypomethylation, genes were identified only in the analyses focusing on transcriptionally active regions in both the SLE and MS datasets in which disease-associated genes were detected. In SLE we identified the involvement of multiple genes associated with the well-established type I interferon pathway, which was not detected in the original EWAS. In MS, interferon-related pathways were likewise detected; this is likely attributable to the fact that 65% of the patients included in the MS EWAS were receiving treatment, and 69.2% of those treated were on interferon therapy ^59^. Focusing on active chromatin regions and leveraging eQTM-based identifications for disease-associated genes are therefore informative. Accordingly, in Table S19 we provide, for each CpG on the methylation array, the sequence context (e.g., solo-WCGW), chromatin state annotation (active or repressed), and the eQTM information detected in the study by Keshawarz *et al.* ^41^ (i.e., the genes with which it is positively or negatively correlated, the number of CpGs correlated with that gene, and the rank of the CpG in terms of association strength within that gene).

Finally, one of the most important observations in this study is that methylation levels at solo-WCGW sites in repressed chromatin were highly consistent within individuals, allowing their average level to be summarized as a hypomethylation index. We hypothesized that if immune activation promotes cell proliferation, the degree of such proliferation might be reflected in this index. Based on this hypothesis, we examined whether methylation changes in transcriptionally active regions correlate with the hypomethylation index. In the NT1 analysis of CD4⁺ T cells, a DMR within the TRA locus was identified as significantly associated with the index, whereas in the MS analysis, a DMR within the IGH locus, which encodes the B-cell receptor heavy chain, showed a similar association after adjustment for cell composition. Notably, in NT1, samples with lower values of the index—indicating more extensive hypomethylation—showed increased T-cell receptor clonality, suggesting expansion of specific T-cell clones. Meanwhile, in B cells from MS patients, hypomethylation has been reported, with lower methylation levels associated with higher IgM levels ^60^. This observation supports the BCR-related signal suggested by our MS analysis. Together, these findings suggest that the hypomethylation index may function as an epigenetic footprint of cumulative immune activation. Although variation in immune cell composition may influence DNA methylation patterns, our observations were consistent across multiple immune-mediated diseases and remained evident after adjustment for cell composition, suggesting that the observed hypomethylation largely reflects proliferation-associated processes rather than differences in cell-type proportions.

This study has two major limitations. First, for diseases other than NT1, the methylation data were derived from whole-blood DNA, which prevented the evaluation of each immune cell subset individually. In the case of NT1, global hypomethylation in repressed chromatin was observed in both CD4^+^ and CD8^+^ T cells, and a similar trend was observed when assessing whole blood. Therefore, unless the proliferating cells constitute a markedly small fraction of the immune cell subset, the impact could be reasonably evaluated even using whole blood. Second, methylation levels were determined using arrays in all of the EWASs we examined. Methylation arrays, particularly the 450K array, are designed to prioritize the analysis of methylation regions upstream of genes that may affect gene expression. Therefore, obtaining deeper coverage using whole-genome bisulfite sequencing could potentially expand our understanding of methylation in regions such as repressed chromatin.

In conclusion, we found that some of the hypomethylated CpGs reported in EWASs of immune-mediated diseases are affected by enhanced proliferation of immune cells, leading to passive demethylation of repressed chromatin, particularly in the solo-WCGW context (Fig 5). This hypomethylation may represent an epigenetic footprint of cumulative immune activation and could serve as a molecular proxy for cumulative immune burden during disease development. By focusing on CpGs in which active demethylation occurs (thereby excluding the effects of random methylation loss), we were able to identify more disease-relevant pathways. These findings provide a conceptual framework for interpreting disease-associated methylation changes and may help refine EWAS strategies for diseases involving cell proliferation.

## Materials and Methods

Extended descriptions of the methods are provided in the Supplementary Methods.

### EWAS data for immune and non-immune diseases

For NT1, we used an in-house dataset (see Supplementary Methods for details). After identifying significant hypomethylation of solo-WCGW in NT1, we examined whether a similar pattern occurs in other immune-mediated diseases. We focused on studies that provide comprehensive data on disease-associated methylation sites in EWASs of other immune-mediated diseases. To further enhance the reliability of the results, we selected studies with sample sizes of >100 per group (cases and controls), including research on SLE ^26^, RA ^27^, MS ^28^, MS severity ^29^, atopic asthma ^30^, and non-atopic asthma ^30^ (Table S1). Additionally, we analyzed EWAS datasets on non-immune-mediated diseases meeting similar criteria as references, focusing on schizophrenia ^31^, pulmonary arterial hypertension ^32^, and Alzheimer’s disease ^33^ (Table S2).

For the disease-associated methylation sites, we used data from studies that provided comprehensive information regarding these sites. However, for the MS study, no information on comprehensive disease-associated methylation sites was available, and only raw data were provided; therefore, we conducted our own data analysis (see Supplementary Methods for details).

### Analysis of solo-WCGW

As all EWASs analyzed in this study used Illumina methylation arrays (450K or EPIC), we summarized the sequence context surrounding each CpG site based on the flanking sequence information provided in the Manifest files supplied by Illumina using a custom Python script (CpGContextCounter.py). Following previous studies ^19,21^, CpG sites without other CpGs within 35 bp upstream or downstream were defined as *solo-CpGs*. CpG sites flanked by A or T (denoted as ‘W’) on both sides were defined as *WCGW*. We assessed whether each disease-associated hypomethylated CpG site corresponded to a solo-WCGW site. We compared the proportion of solo-WCGW sites among hypomethylated DMPs in each disease with that among all analyzed CpGs on the corresponding methylation platform (450K or EPIC) used for each disease (Fig S8). Statistical analyses, including the calculation of *P*-values using Fisher’s exact test and the estimation of ORs, were performed using R software.

### Analysis of PMDs

To investigate the impact of PMDs in non-cancerous immune cells, we identified PMDs using whole-genome bisulfite sequencing data ^35^ from immune cells with, MethPipe ^61,62^. In addition to the 15 immune cell subsets listed in Table S3, we included neurons as a reference for non-cancerous, non-immune cells. The nine whole-genome bisulfite sequence data of tumor samples ^19^ were also included as a representative for cancerous cells. For each cell type, PMDs were individually delineated, and PMDs detected in each cell type were consolidated, such that any region identified as a PMD in at least one cell was considered a PMD. Based on this criterion, PMDs were compiled separately for immune cells, neurons, and cancer cells. PMD detection, calculation of the total span of detected regions, assessment of overlapping regions, and identification of EPIC/450K array probes located within the detected PMDs were performed using PMD_processing.sh (Additional_file_2), region_length_summary.py, and pairwise_overlap_length.py, respectively. Principal component analysis was performed using R software. To investigate the chromatin states enriched in PMD regions, we utilized ATAC-seq, DNase-seq, and histone ChIP-seq data from the ENCODE project ^63^. The detailed parameters for data processing for immune cells are described in the “Examination of regions characterized by chromatin accessibility and histone modifications” section. The analyses for neuron and tumor samples were consistent with those for immune cells, except for the tissue types, which were determined by the biosample metadata. The list of biosamples used for neuron and tumor samples is provided in Table S20. Due to the lack of available data for H3K9me2 in the tumor and neuronal analyses and H3K9ac in the neuron analysis, we substituted blood-derived data for these histone marks. We compared the proportion of solo-WCGW sites among hypomethylated solo-WCGW sites in PMD with all analyzed solo-WCGW sites (Fig S8).

### Examination of chromatin accessibility, histone modifications, and replication timing

To investigate in more detail the characteristics of regions containing hypomethylated solo-WCGW CpGs, particularly those associated with chromatin accessibility and histone modifications, we utilized publicly available ATAC-seq, DNase-seq, and histone ChIP-seq datasets from the ENCODE project ^63^. Further details, including replication timing analyses, are described in the Supplementary Methods.

### Investigation of the effect of cytokines on DNA methylation and solo-WCGW

To investigate the impact of cytokines on DNA methylation, we targeted previous studies that compared methylation rates in cells before and after cytokine treatment. GEO datasets (GEO GSE144804, GEO GSE56621) from two previous studies were used, focusing on TGF-β^39^ and TNF-α^38^. For the analysis of TNF-α, human umbilical vein endothelial cells (n = 37) were treated with 20 ng/mL of TNF-α, and the data before and after treatment (24 h) were compared. A *t*-test was performed using the filtered and standardized β-values. Given that the sample size was sufficient, the analysis was based on the threshold *P*<1.0E-04, which is the registration requirement for the EWAS catalog ^64^. The analysis of TGF-β used methylation data acquired before and after treatment (120 h) of ovarian cancer cells with 5 ng/mL of TGF-β. Both a *t*-test and Wilcoxon rank-sum test were conducted because the number of samples in this dataset was small (n = 3). CpG sites were considered associated with TGF-β treatment if they satisfied the criteria of *P*<0.05 in the *t*-test and the theoretical minimum *P*-value threshold in the rank-sum test (*P*<0.1). The associations between the cytokines and solo-WCGW and disease-associated methylation sites were summarized in the same manner as described above. Chromatin accessibility and histone marks were analyzed by setting the ENCODE organ to “endothelial cell” for the TNF-α dataset and “ovary” for the TGF-β dataset. The procedure followed the same steps as described in the Supplementary Methods (Examination of regions characterized by chromatin accessibility and histone modifications), except that no ATAC-seq data for endothelial cells were available in the ENCODE database.

### Definitions of active and repressed chromatin regions

In this study, genomic regions were classified as those with open chromatin and active transcription, and those that are repressed. Following previous studies, the regions were defined based on a combination of detection by ATAC-seq or DNase-seq and the presence of annotated histone marks. Specifically, regions detected by either ATAC-seq or DNase-seq and marked with at least two of three histone modifications (H3K4me3, H3K9ac, or H3K27ac) were considered active promoters, whereas those marked with histone modifications H3K4me1 and H3K27ac were considered active enhancers ^65^. The union of these regions was designated as an “active region”. Genomic regions that did not fall into any of these categories were defined as “repressed chromatin”.

### Distribution of CpGs in mQTLs and eQTMs across active and repressed chromatin regions

We examined the relationship between transcriptionally active/repressed regions and CpGs located in mQTLs or eQTMs. For mQTLs, we used loci identified in previous large-scale studies that measured DNA methylation in whole blood using the Illumina EPIC array ^40^. For eQTMs, we used data from two studies registered in the eQTM-Catalog (https://shiny.crc.pitt.edu/eqtm_browser/), both of which measured DNA methylation in whole blood ^41,42^. Although one study was profiled on the EPIC array, its sample size was <50 ^42^; therefore, we complemented it by analyzing eQTM data identified in more than 2,000 samples using the 450K array ^41^. Among CpGs assayed on the EPIC array, we compared the proportion of mQTL versus non-mQTL CpGs located in active regions. A similar comparison was performed for eQTMs, focusing on CpG sites measurable with the Illumina 450K array. We also assessed CpGs shared between mQTLs and eQTMs. For eQTMs, analyses were conducted both using the union of eQTMs detected in either study and using only those detected in both studies.

### Identification of DMRs associated with the hypomethylation index

To assess whether hypomethylation at solo-WCGW sites in repressed chromatin showed consistent inter-individual variation across samples, we visualized methylation levels using heatmaps generated in R. Because the relative levels of hypomethylation were largely consistent within individuals across these sites, we summarized their average methylation level as a hypomethylation index.

We then searched for DMRs in transcriptionally active regions whose methylation levels were associated with this hypomethylation index using the bumphunting method implemented in the R package minfi ^66^. In the NT1 CD4⁺ T-cell analysis, the ratio of naïve CD4⁺ T cells (naïve CD4/CD4) estimated using the DNA Methylation Age Calculator, together with age, sex, and disease status, was included as covariates. In the NT1 CD8⁺ T-cell analysis, the ratio of naïve CD8⁺ T cells (naïve CD8/CD8), age, sex, and disease status were included as covariates. In the MS analysis, estimated immune cell proportions (CD8, NK, B cell, monocyte, granulocyte, naïve CD4, and naïve CD8) selected to minimize multicollinearity, along with age, sex, and disease status, were included as covariates.

Only DMRs in which more than 70% of the CpGs were located within transcriptionally active regions, as defined in this study, were retained for further analysis. To ensure the robustness of the analysis, probes located within the *HLA* region (chr6:27,000,000–34,000,000, hg19) were excluded. Genes linked to CpG sites within each DMR through eQTM relationships were assigned to the corresponding DMRs based on previously reported eQTM datasets ^41^.

### Estimation of T-cell receptor clonality

For NT1, T-cell receptor (TCR) clonality was estimated using whole-blood RNA-seq data derived from the same samples used in the DNA methylation analysis. Details of the RNA-seq data have been described previously ^25^. TCR repertoires were reconstructed using TRUST4 ^43^. Only productive TCR sequences were included in the analysis. Clonotypes were defined based on CDR3 nucleotide sequences, following the approach described in the original TRUST4 study. Clonality was calculated as 1 − normalized Shannon entropy of clonotype frequencies. Clonality metrics were calculated separately for the TRA and TRB repertoires.

### Prioritizing disease-associated genes through eQTM mapping

To prioritize disease-associated genes from disease-related hypomethylated loci, we utilized eQTM relationships ^41^. Detailed methods are described in the Supplementary Methods.

### Bioinformatics workflow and command lines

The command lines used in this study are listed in Additional file 2. The Python programs are also available on GitHub (https://github.com/mihshimada/Hypomethylation_immune).

## Supporting information

Command lines

## Acknowledgements

We thank all participants who cooperated in this study. We also thank the staff of Seiwa Hospital and Koishikawa Tokyo Hospital for their support in blood sampling.

## Declarations

### Ethics approval and consent to participate

Written informed consent was obtained from all participants or their families, and the protocols were approved by the ethics committees of all collaborating institutes.

### Availability of data and materials

NT1 EWAS data has been deposited in the DNA Data Bank of Japan (DDBJ; https://www.ddbj.nig.ac.jp/index-e.html), and is available from the European Nucleotide Archive operated by the European Molecular Biology Laboratory’s European Bioinformatics Institute (EMBL-EBI; https://www.ebi.ac.uk/ena/browser/home) under the accession number PRJDB15980.

The command lines used in this study are listed in Additional file 2. The python programs are also available on GitHub (https://github.com/mihshimada/Hypomethylation_immune).

Any additional information required to reanalyze the data reported in this paper is available from the lead contact upon request.

### Authors’ contributions

MS and YO designed the study. MH collected NT1 samples. MS, TM, and YH carried out experiments and statistical analyses. YH, TM, MH, TK, and KT interpreted the results. MS and YO drafted the manuscript, and all authors contributed to the final version of the paper.

