## Supplementary methods and figures for "Replication-associated solo-WCGW hypomethylation reflects cumulative immune activation across diseases"

**Supplementary Information (Additional file 1)**

### Supplementary Methods

#### EWAS data for CD4$^+$ and CD8$^+$ T cells from NT1

In this study, we first focused on NT1, which is the primary subject of our research. Briefly, 42 (discovery: 28, replication: 14) patients with NT1 and 42 (discovery: 28, replication: 14) control subjects without significant sleep-disturbing events who underwent diagnostic sleep studies (polysomnogram and multiple sleep latency test) were included. NT1 was diagnosed by sleep specialists according to the International Classification of Sleep Disorders, 3rd edition [1]. CD4$^+$ and CD8$^+$ T cells were extracted from the lymphocyte fraction, and DNA was extracted from the cells. DNA methylation levels were examined using Infinium® Methylation EPIC BeadChip (Illumina, San Diego, CA, USA). The methylation rate was calculated as the β-value, and filtering and normalization (BMIQ [2], quantile normalization, and Combat [3]) were performed. M-values were used in the case-control analysis. As there were no significant differences between cases and controls in terms of background factors such as age and sex, disease-associated differentially methylated sites were identified using a *t*-test. Details regarding the EWAS data are described elsewhere [4].

#### Whole-blood EWAS data in NT1

The DNA methylation data obtained using the Infinium HumanMethylation27 BeadChip (Illumina) on whole blood-derived DNA from individuals with NT1 were reported in our previous study [5].

#### Pipeline for MS dataset

For the MS study, as only raw data were provided, we conducted our own data analysis.

Using β-values normalized according to the default settings of the R package minfi [6], we estimated the cellular composition ratios using the DNA Methylation Age Calculator [7]. Because significant differences in the ratios of NK cells and $CD8^+$ T cells ($P$ = 2.21E-08 and $P$ = 0.00702) were observed between patients and healthy controls, the case-control analysis was conducted using the estimated cellular composition ratios, age, and sex as covariates. To assess the overall impact, CpG sites exhibiting $P<0.01$ were considered disease-associated methylation sites.

**Examination of regions characterized by chromatin accessibility and histone modifications**

To investigate in more detail the characteristics of regions containing hypomethylated solo-WCGW CpGs, particularly those associated with chromatin accessibility and histone modifications, we utilized publicly available ATAC-seq, DNase-seq, and histone ChIP-seq datasets from the ENCODE project [8]. We used all datasets with an assay title of Histone ChIP-seq, ATAC-seq, or DNase-seq that satisfied the following conditions: Organism: Homo sapiens; Organ: blood; Perturbation: not perturbed. Only datasets aligned to the GRCh38 genome assembly and available in BED format (narrowPeak) were considered. Furthermore, to ensure data quality, only datasets free from any flagged issues in the ENCODE audit category were included in the analysis. The retrieved data were merged by assay type, and in the case of Histone ChIP-seq, the data were separately merged for each histone mark. Genomic regions that appeared in at least one dataset of interest were considered associated regions and included in the analysis, as implemented in the script mergeBEDbyTarget.py. The overlap between hypomethylated solo-WCGW in each disease, and the regions identified by each assay, were assessed using the script

GenomicFeatureLocator.py. We compared the proportion of solo-WCGW sites among hypomethylated solo-WCGW sites in repressed regions with all analyzed solo-WCGW sites (Fig S8).

**Investigation of replication timing**

To investigate the replication timing of each gene region in non-cancerous immune cells, we explored the ENCODE Replication Timing Series [8], focusing on *Homo sapiens*, "Blood" as the tissue type, and non-cancerous cells. Data from samples GM06990, GM12878 (adult), GM12801, and GM12812 were used in this analysis. The percentage normalized signal data in BigWig format for the G1b, G2, S1, S2, S3, and S4 phases were converted to BedGraph format and subsequently merged using merge_bedgraph.sh. G1, S1, and S2 were treated as early replication phases, whereas S3, S4, and G2 were treated as late replication phases. Replication timing signals were analyzed based on the method described by Dietzen et al. [9]. Quantile normalization of $\log_2$(early/late) was performed using the RT signal of GM06990 as a reference, with the normalize.quantiles.use.target function (preprocessCore v1.44) in R software. To reduce signal noise, LOESS smoothing was applied to each chromosome using the 'loess' function in R software, with a span of 300 kb. The smoothed values were used as the final RT signal, where positive values represented early-replicating regions, and negative values represented late-replicating regions. The same analysis was performed for four samples, and regions consistently classified as early or late in all samples were defined as early or late, respectively. Regions that did not show consistent classification across all four samples were defined as non-conserved. Finally, the CpG sites targeted by the EPIC and 450K arrays were assigned to early, late, or non-conserved replication timing regions using the annotate_probes.sh script. Based on this classification, we compared the proportion of solo-WCGW sites among hypomethylated solo-WCGW sites in late-replicating regions with all analyzed solo-WCGW sites (Fig S8). Statistical tests were performed, following the same procedures as described in the “Analysis of solo-WCGW” section.

### Prioritizing disease-associated genes through eQTM mapping

To prioritize disease-associated genes from disease-related hypomethylated loci, we utilized eQTM relationships. In this analysis, we focused on cis-eQTMs identified at $P < 1\times10^{-7}$ in the large-sample study by Keshawarz *et al* [10]. Specifically, we examined genes whose expression was associated, as eQTMs, with hypomethylated disease-associated CpG sites residing in transcriptionally active and repressed regions. Notably, multiple CpG sites often correlate (as eQTMs) with a single gene, with varying strengths of association. Because calling a gene disease-associated when, for instance, only the weakest of those CpGs is disease-related is prone to false positives, we employed the gene-selection procedure depicted in Fig S7. First, for each gene, we considered all CpG sites that formed an eQTM with that gene and ranked them by the strength of association. We then designated a gene as a candidate disease-associated gene if it satisfied any of the following criteria: (1) the top-ranked CpG for that gene was disease-associated; (2) ≥70% of the eQTM-linked CpGs for that gene were disease-associated; or (3) ≥50% of the eQTM-linked CpGs were disease-associated and, among them, at least one CpG ranked within the top five by association strength. In addition, to further reduce false positives, we required that each candidate be supported by eQTM evidence from ≥3 distinct CpG sites. The mapping of each disease-associated hypomethylated CpG to its eQTM-linked gene and the ranking of that CpG among all CpGs forming eQTMs with the gene (by strength of association) were computed using eQTMgeneRanker.py. In addition, for CpG sites on the Illumina 450K array that have eQTM-linked genes, the aggregation of associated genes and the within-gene rank of each CpG among all eQTM CpGs was performed with eQTMprobeAggregator.py.

For SLE and MS, in which multiple candidate disease-associated genes were identified, we further assessed whether these genes show disease-associated expression in transcriptomic datasets [11-13]. For SLE whole-blood expression, we analyzed GSE65391 and, following the methodology used in the original study, identified SLE-associated genes by fitting a linear mixed-effects model with age as a covariate and subject-specific random intercepts to accommodate repeated measures, followed by multiple-testing correction using the Benjamini–Hochberg procedure. In addition, for SLE, we referred to another study that profiled gene expression in fractionated immune cell subsets [12]; as complete lists of associated genes were publicly available, we used these as reference. The same procedure was applied to the MS dataset [13].

### Protein-protein interaction analysis using STRING

To explore the characteristics of genes implicated in disease by eQTM-based analysis, we performed a protein-protein interaction analysis using STRING [14]. The STRING analysis was performed using default settings, with the minimum required interaction score set to medium confidence (0.400). We performed k-means clustering based on the number of protein-protein interaction groups.

### Supplementary Figures

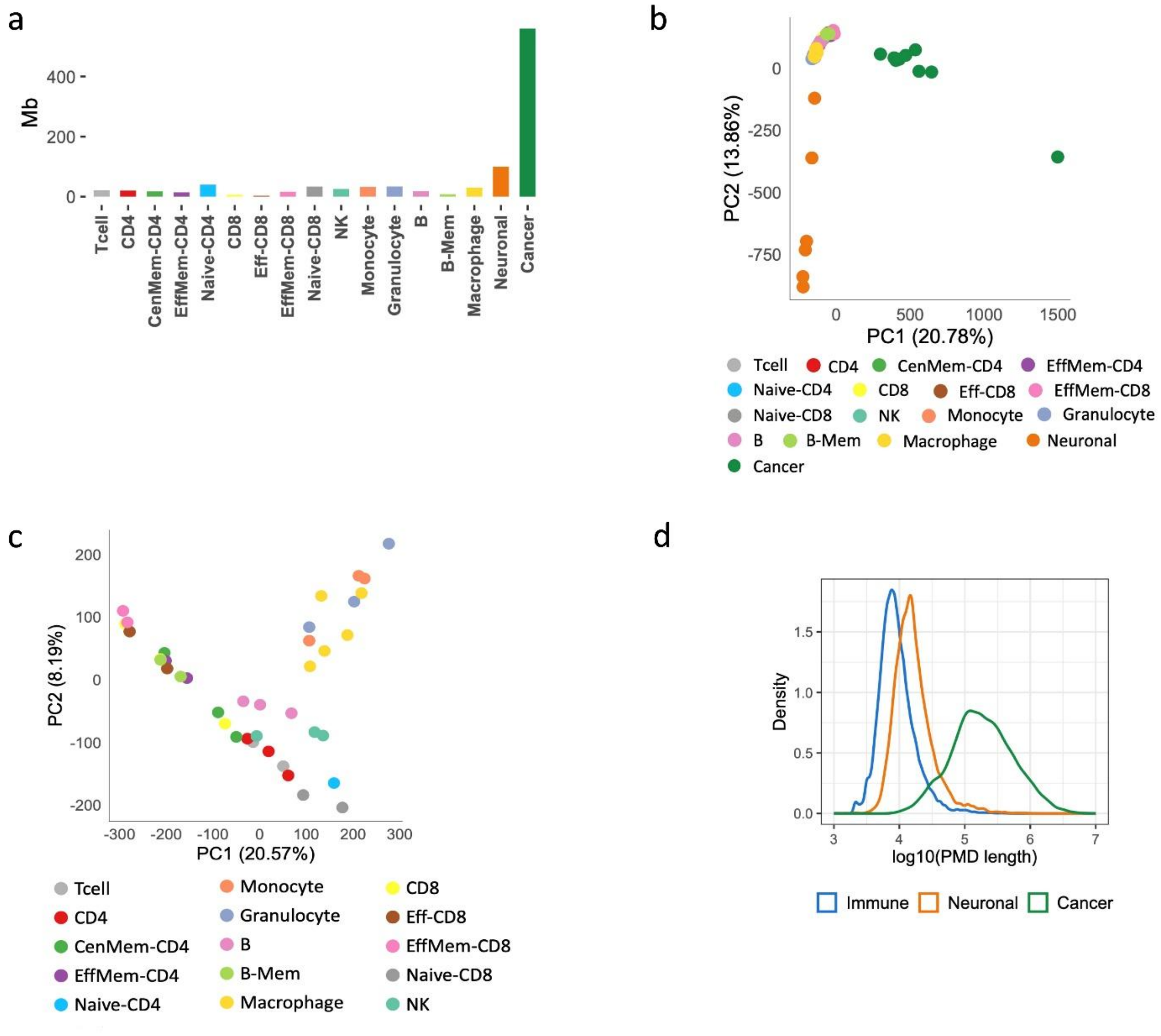


**Fig. S1. PMD detection in normal and cancer cells. a.** Size (Mb) of regions identified as PMDs in each cell type or tissue. **b.** Principal component analysis (PCA) based on CpG sites located within PMDs identified in each cell type or tissue. **c.** PCA restricted to immune cell subtypes. PC1 clearly separates effector memory cells from naïve cells and reveals clustering according to immune cell subsets. **d.** Distribution of PMD lengths.

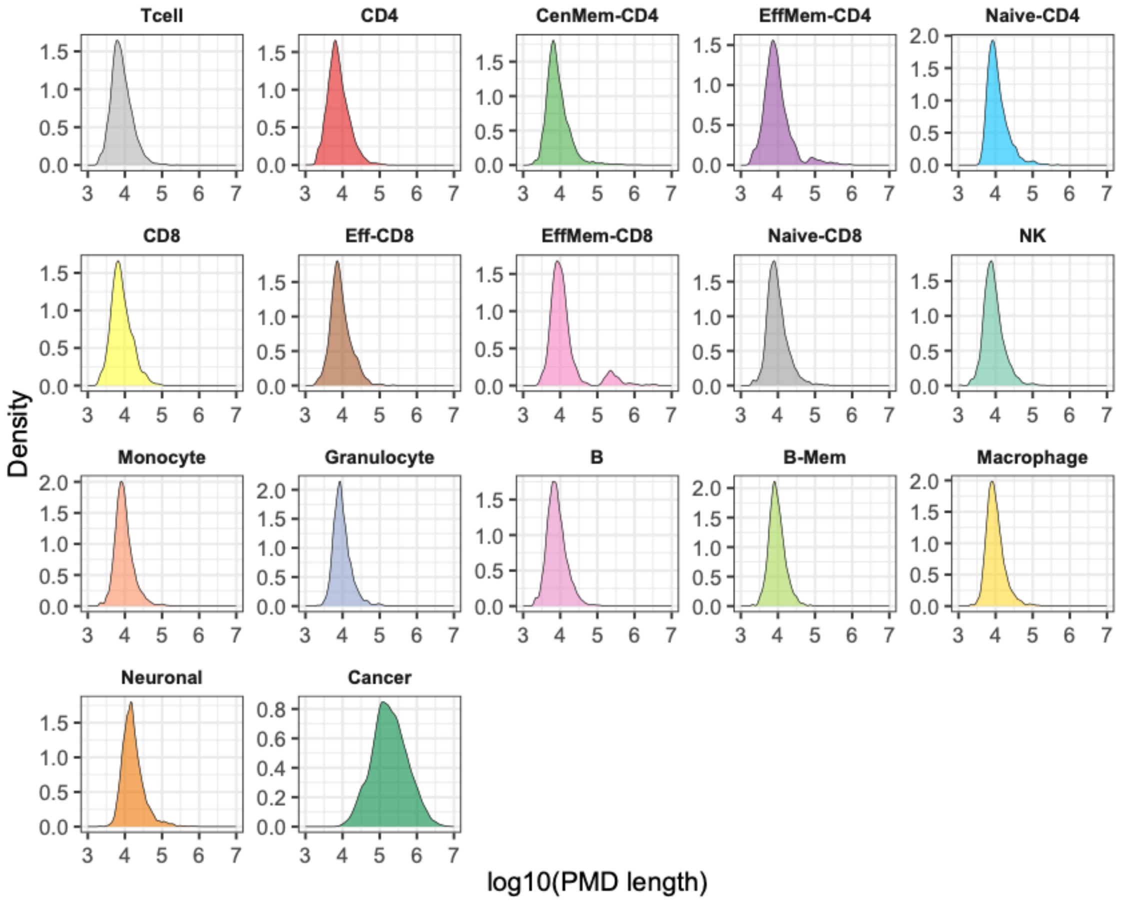


**Fig. S2. Distribution of PMD length across cell and tissue types.**

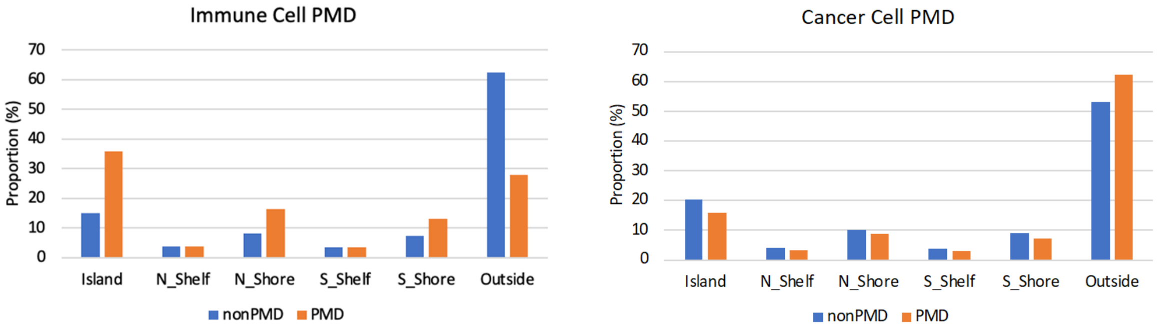


**Fig. S3. Proportion of CpG sites covered by the EPIC array, located within PMDs identified in immune cells and cancer cells, distributed across CpG islands, shores, shelves, and outside regions.**

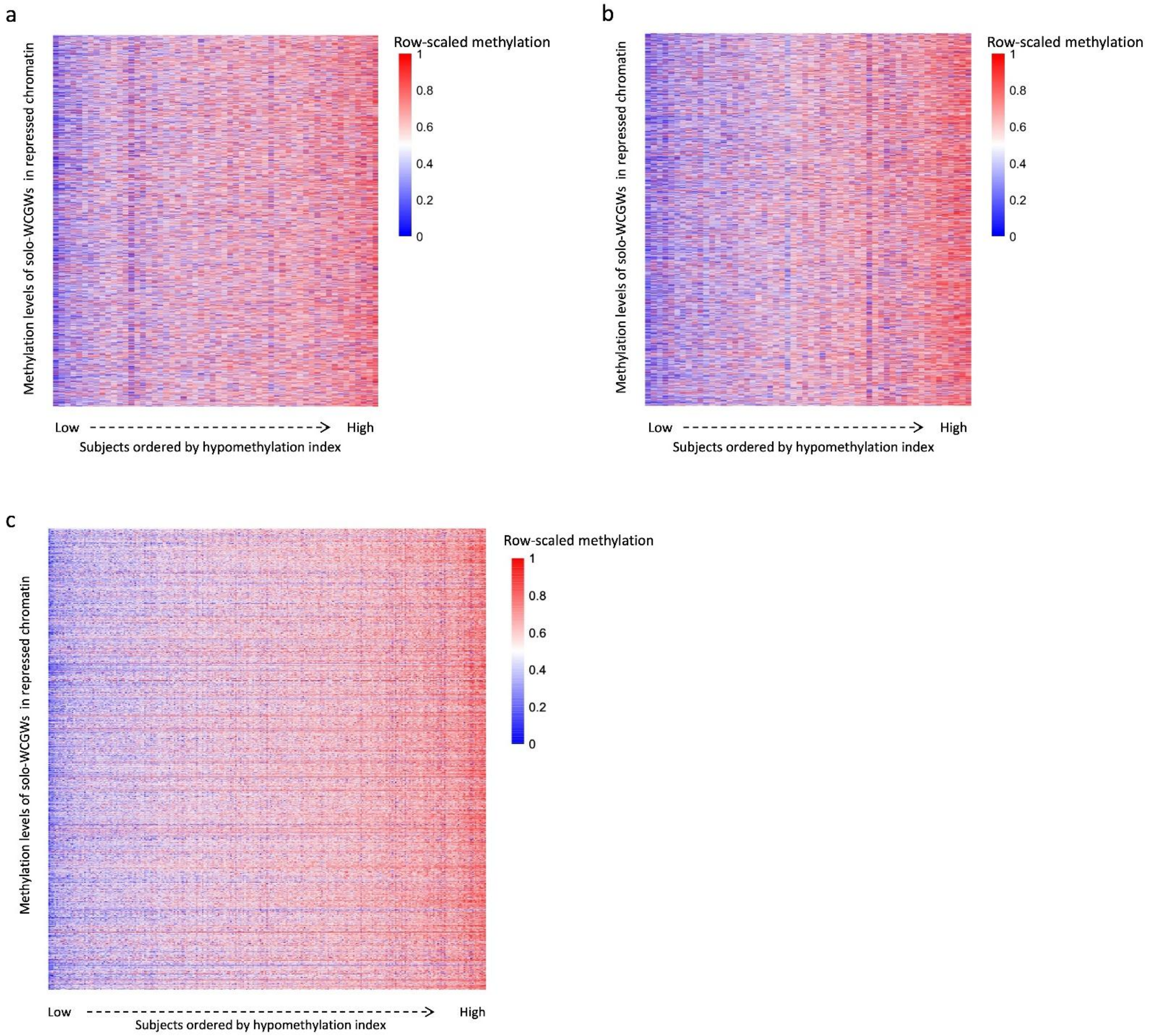


**Fig. S4. Consistent inter-individual variation in methylation levels at solo-WCGW sites in repressed chromatin.** Heatmaps showing row-scaled methylation levels of solo-WCGW CpG sites located in repressed chromatin. Rows represent CpG sites and columns represent individual samples ordered by the hypomethylation index. Methylation levels show consistent inter-individual variation across these sites, supporting the use of their average methylation level as a hypomethylation index. **a.** NT1 $CD4^{+}$ T cells. **b.** NT1 $CD8^{+}$ T cells. **c.** Multiple sclerosis whole-blood samples.

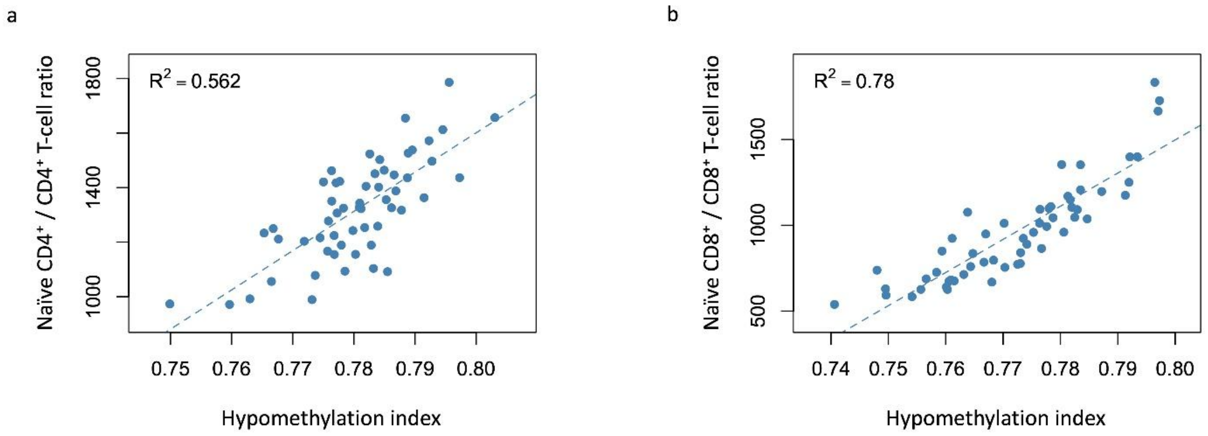


**Fig. S5. Association between the hypomethylation index and estimated naïve T-cell ratios in NT1.** Scatter plots showing the relationship between the hypomethylation index and methylation-based estimates of naïve T-cell ratios in **a.** $CD4^+$ T cells and **b.** $CD8^+$ T cells. Dashed lines indicate linear regression fits. The naïve $CD4^+/CD4^+$ and naïve $CD8^+/CD8^+$ values were derived from the DNA Methylation Age Calculator and should be interpreted as relative methylation-based estimates rather than direct measurements of cell proportions.

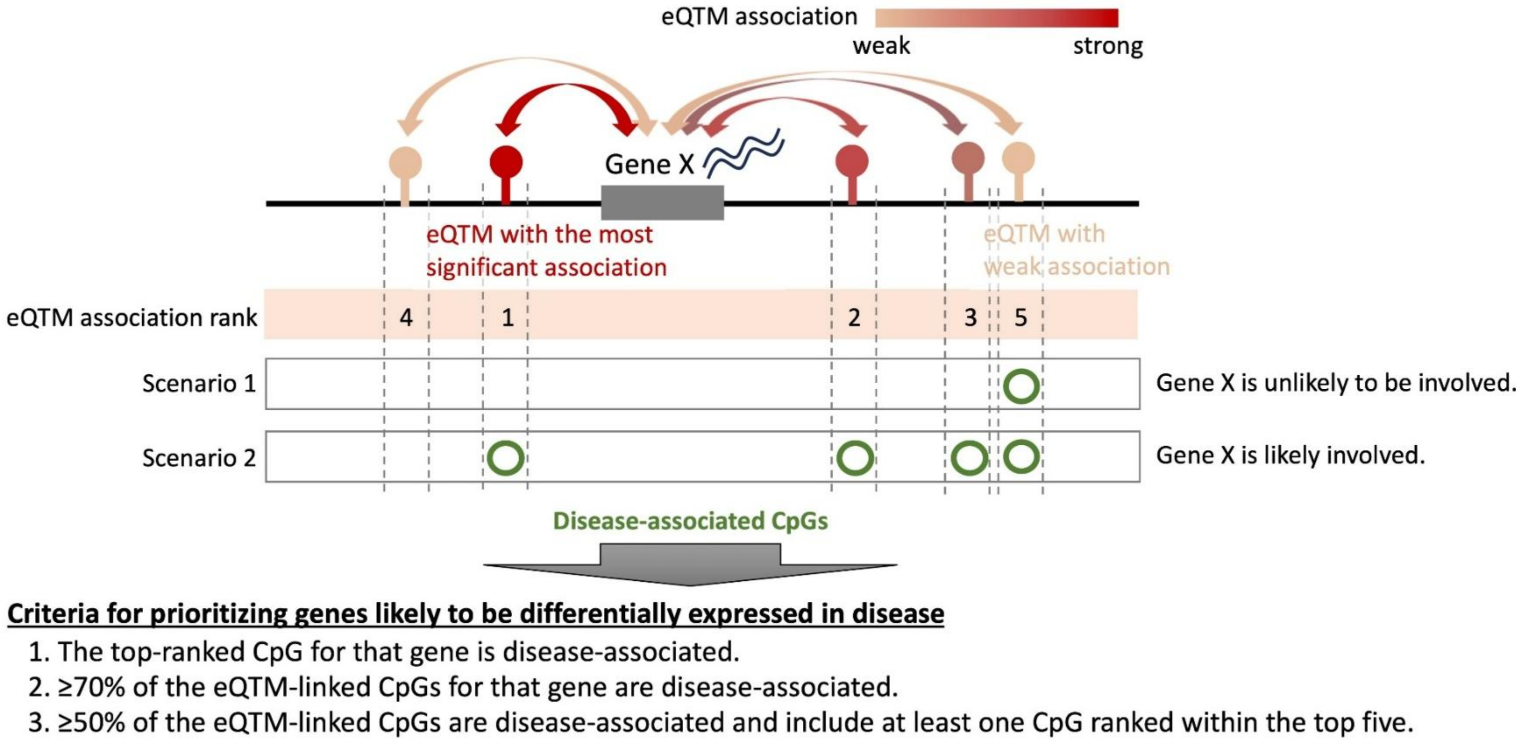


**Fig. S6. Exploring disease-associated genes using eQTM relationships.** Many genes are linked, via eQTMs, to multiple methylation sites. In such cases, if only the methylation site belonging to the weakest eQTM is disease-associated, it is unlikely that the corresponding gene is truly differentially expressed in the disease (Scenario 1). By contrast, if many of the eQTMs for a given gene are disease-associated, or if the methylation site in the strongest eQTM is disease-associated, the probability that the gene's expression is altered in the disease is higher (Scenario 2). Based on this rationale, we defined criteria to identify more reliable candidate genes with disease-related expression changes.

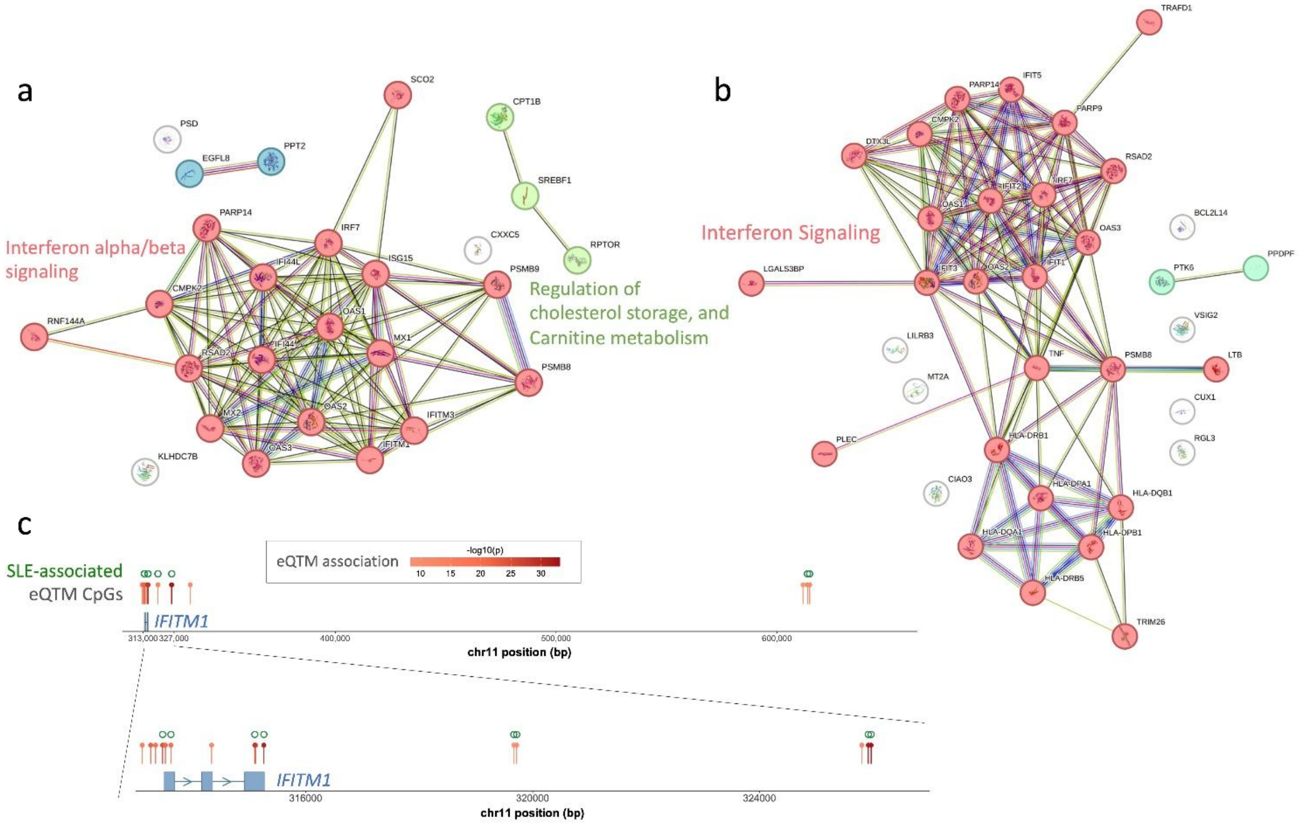


**Fig. S7. EWAS analysis integrating eQTM relationships in transcriptionally active regions. a.** Genes predicted to be upregulated in SLE based on disease-associated hypomethylated CpG sites and their eQTM relationships. **b.** Genes predicted to be upregulated in MS. **c.** Example of multiple CpG sites showing eQTM relationships: the *IFITM1* locus in SLE.

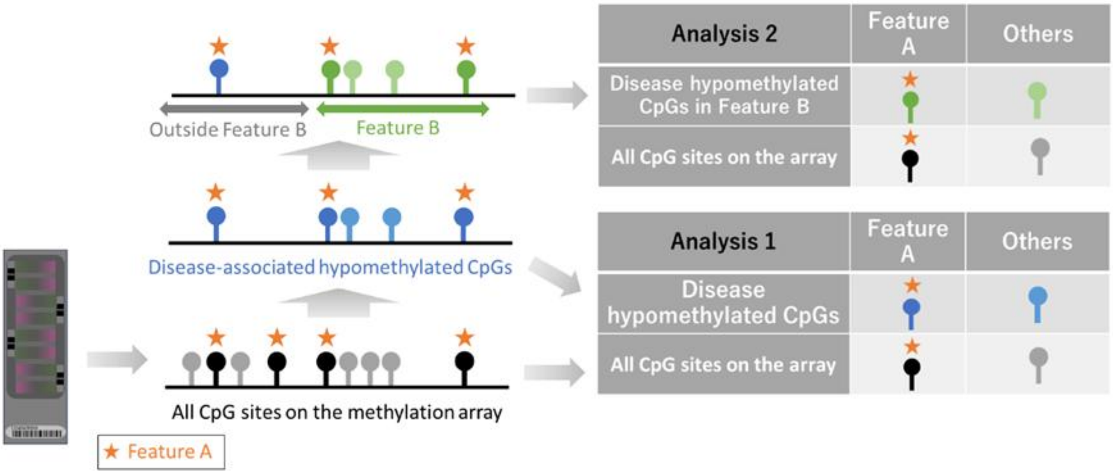


**Fig. S8. Calculation method of ORs.** This figure provides a conceptual illustration of how the ORs shown in Fig. 1 and Fig. 2e were calculated. In Fig. 1, ORs were derived using the Analysis 1 approach. In this analysis, “Feature A” represents “solo-“, “WCGW-“, or “solo-WCGW” CpGs. The proportion of Feature A among disease-associated hypomethylated CpGs was compared to its proportion among all analyzed CpG sites on the array. In Analysis 2 (Fig. 2e), we further focused the analysis on disease-associated hypomethylated sites overlapping with Feature B. Here, Feature B refers to immune-cell PMDs, late-replicating regions, or repressed chromatin regions.
