## Supplementary material for "Replication-associated solo-WCGW hypomethylation reflects cumulative immune activation across diseases": Command lines

**Command lines used in this study:**

These code snippets represent the commands used in the study but are simplified for clarity and reproducibility.

***PMD detection using methpipe (PMD_processing.sh)***

\# This script performs PMD analysis using methpipe for a list of .meth files.

\# Usage:

\#	1. Prepare list.txt with one .meth file path per line.

\#	2. Edit the conda environment and paths below as needed.

#!/bin/bash

#------------------------ Activate the conda environment

\# Edit the path below to match your local conda installation and environment name

source /path/to/miniconda3/bin/activate your_env_name

#------------------------ Set environment variables

\# Replace the following paths with your own paths if necessary

export LD_LIBRARY_PATH=/path/to/miniconda3/envs/your_env_name/lib:$LD_LIBRARY_PATH

export PATH=/path/to/methpipe/methpipe-5.0.1/bin:$PATH

#------------------------ Process each file listed in list.txt

while IFS= read -r file; do

output_file="${file}_result.txt"

> "$output_file" # Initialize output file

for chr in {1..22}; do

\# Extract lines for the current chromosome from the .meth file

awk -v chr="chr$chr" '$1 == chr' "$file" > "chr${chr}.meth"

\# Run PMD analysis

echo "Running PMD for chr$chr"

pmd -i 1000 -o "chr${chr}.pmd" "chr${chr}.meth"

\# Append PMD results to the final output

cat "chr${chr}.pmd" >> "$output_file"

\# Clean up temporary files

rm -f "chr${chr}.meth" "chr${chr}.pmd"

done

echo "Processed $file -> $output_file"

done < list.txt

***R code***

#------------------------ Enrichment Analysis Script

############################################

\# README: Enrichment Analysis Script

\# Description:

\# This script reads a table of counts and performs

\# proportion and chi-square and Fisher's exact tests for each row.

\# It calculates p-values, odds ratios, and confidence intervals.

\#

\# Input file:

\# - input.txt (tab-delimited, with appropriate column headers)

\#	This file should be placed in the same directory

\#	where the script is executed.

\#	The input file should be a tab-delimited `.txt` file with a header row.

\#	Each row represents a test item (e.g., transcription factor),

\#	and the columns should include counts for control and case groups.

\#	ID	Ctrl_Pos	Ctrl_Neg	Case_Pos	Case_Neg

\#	TF1	10	90	20	80

\#	TF2	15	85	30	70

\#	...

\#	**Note**: The order of columns (control vs. case) must be correctly specified.

\#

\# Output file:

\# - result_summary.csv

############################################

\# Read input data

data <- read.table("input.txt", header = TRUE)

output_file <- "result_summary.csv"

\# Prepare an empty list to store results

results <- list()

\# Loop through each row

for (i in 1:nrow(data)) {

\# Extract ID or label (e.g., TF name, region ID, etc.)

row_id <- as.character(data[i, 1]) # 1st column

\# Extract contingency values and convert to numeric

group_control <- as.numeric(data[i, 2:3]) # Control group

group_case <- as.numeric(data[i, 4:5])	# Case group

if (group_control[1] > 0 || group_case[1] > 0) {

\# Create 2x2 matrix

contingency_matrix <- matrix(c(group_case, group_control), ncol = 2, byrow = TRUE)

\# Chi-squared test

prop_test_result <- prop.test(contingency_matrix)

chi_statistic <- prop_test_result$statistic

p_chi <- prop_test_result$p.value

\# Fisher's exact test

fisher_result <- fisher.test(contingency_matrix)

p_fisher <- fisher_result$p.value

\# Proportions

rate_control <- group_control[1] / sum(group_control)

rate_case <- group_case[1] / sum(group_case)

\# Odds ratio and confidence interval

odds_ratio <- (contingency_matrix[1,1] / contingency_matrix[1,2]) /

(contingency_matrix[2,1] / contingency_matrix[2,2])

log_or <- log(odds_ratio)

se <- sqrt(sum(1 / contingency_matrix))

conf_interval <- exp(log_or + qnorm(c(0.025, 0.975)) * se)

\# Store result

results[[i]] <- data.frame(

ID = row_id,

rate_case = rate_case,

rate_control = rate_control,

chi_squared = chi_statistic,

p_chi = p_chi,

p_fisher = p_fisher,

odds_ratio = odds_ratio,

CI_lower = conf_interval[1],

CI_upper = conf_interval[2],

case_positive = group_case[1],

case_negative = group_case[2],

control_positive = group_control[1],

control_negative = group_control[2]

)

} else {

\# For rows with no signal

results[[i]] <- data.frame(

ID = row_id,

rate_case = NA,

rate_control = NA,

chi_squared = NA,

p_chi = NA,

p_fisher = NA,

odds_ratio = NA,

CI_lower = NA,

CI_upper = NA,

case_positive = group_case[1],

case_negative = group_case[2],

control_positive = group_control[1],

control_negative = group_control[2]

)

}

}

\# Combine all into one data frame

results_df <- do.call(rbind, results)

\# Write with headers to CSV

write.csv(results_df, output_file, row.names = FALSE, quote = FALSE)

#------------------------ Forest plot

\# This script generates a forest plot using odds ratios and confidence intervals.

\# It assumes the input data includes groupings for color differentiation.

\# Requires the ggplot2 package.

\# Load ggplot2 library

library(ggplot2)

\# Read data file (tab-delimited with header)

\# The data file should have the following columns:

\# - NO: Numeric ID or row index

\# - OR: Odds ratio

\# - lower: Lower bound of 95% confidence interval

\# - upper: Upper bound of 95% confidence interval

\# - group: Group assignment for coloring (e.g., "control" or "case")

DATA <- read.table("data.txt", header = TRUE)

\# Create the base forest plot

re <- ggplot(DATA, aes(x = NO, y = OR, colour = group)) +

geom_point(shape = 20, size = 3) + # Add points

＃scale_y_log10() +	# Log10 scale for odds ratios if needed

theme_linedraw() +	# Clean theme

scale_color_manual(values = c("darkorange2", "blue2")) + # Define colors

geom_errorbar(aes(ymin = lower, ymax = upper), width = 0.07, size = 0.5) + # Confidence intervals

geom_hline(yintercept = 1, linetype = 2, col = "blue") + # Reference line at OR = 1

#scale_x_continuous(breaks = seq(0.5, 29.5, 2)) +	# Customize x-axis if needed

coord_flip() # Flip coordinates for horizontal forest plot

\# Refine plot theme

re <- re + theme(

panel.grid.major = element_line(size = 0.2, colour = "gray"),

strip.placement = "outside",

panel.grid.minor.y = element_blank(), # Remove minor grid lines

panel.grid.minor.x = element_blank(),

strip.text.y = element_text(angle = 0)

)

\# Display the plot

re

#------------------------ Quantile normalization using reference distribution

############################################

\# This script performs quantile normalization on the "Log2_Ratio" column

\# across multiple CSV files using a reference distribution.

\# It uses the preprocessCore package.

\# Input file format:

\# 　　The input CSV file should contain the following columns:

\#

\# 　　Chr	: Chromosome name (e.g., "chr1")

\# 　　Start	: Start coordinate of the genomic window

\# 　　End	: End coordinate of the genomic window

\# 　　Early	: Read counts in early replicating fraction (optional)

\# 　　Late	: Read counts in late replicating fraction (optional)

\# 　　Log2_Ratio : Log2-transformed signal (used for smoothing)

\#

\# 　　Example rows:

\#

\# 　　Chr	Start	End	Early Late Log2_Ratio

\# 　　chr1	24499	26499	55	45	0.289506617

\# 　　chr1	26499	27499	51	49	0.057715498

\# 　　chr1	27499	29499	51	48	0.087462841

############################################

\# Load required package

library(preprocessCore)

\# Set working directory containing the input CSV files

\# (Update this path as appropriate or set externally when sourcing the script)

setwd("your/working/directory/path")

\# Get list of input CSV files that contain "updated_final_result" in their names

file_list <- list.files(

path = ".",

pattern = "updated_final_result.*\\.csv",

full.names = TRUE

)

\# Specify the reference sample file to define the target distribution

reference_file <- "./updated_final_result_GM06990.csv" # Reference sample

reference_data <- read.csv(reference_file)

\# Extract the Log2_Ratio column from the reference, excluding NA values

reference_signal <- reference_data$Log2_Ratio

reference_signal <- reference_signal[!is.na(reference_signal)]

\# Define a function to normalize the Log2_Ratio column using the reference

normalize_log2_ratio <- function(file, reference_signal) {

data <- read.csv(file)

signal <- data$Log2_Ratio

\# Perform quantile normalization only on non-NA values

signal_normalized <- normalize.quantiles.use.target(

as.matrix(signal[!is.na(signal)]),

reference_signal

)

\# Replace the normalized values back into the original data frame

data$Log2_Ratio[!is.na(data$Log2_Ratio)] <- signal_normalized

return(data)

}

\# Apply normalization to all files

normalized_data_list <- lapply(file_list, normalize_log2_ratio, reference_signal)

\# Write normalized data to new CSV files with a "normalized_" prefix

for (i in seq_along(file_list)) {

output_file <- paste0("normalized_", basename(file_list[i]))

write.csv(normalized_data_list[[i]], file = output_file, row.names = FALSE)

}

#------------------------ loess_smoothing

############################################

\# This script applies loess smoothing to the "Log2_Ratio" column

\# in a CSV file that contains replication timing data across the genome.

\# Smoothing is performed separately for each chromosome using a span of 300 kb.

\# The smoothed data is saved as a new CSV file, and an example plot is shown for chromosome 1.

\# Input file format:

\# 　　In the same format as the input file for quantile normalization.

############################################

\# Load the data

data <- read.csv("input_file.csv")

\# Remove rows with NA values

data <- na.omit(data)

\# Get unique chromosomes

chromosomes <- unique(data$Chr)

smoothed_data <- data.frame()

\# Set the smoothing span: 300 kb

span_kb <- 300000

\# Apply loess smoothing for each chromosome

for (chr in chromosomes) {

chr_data <- subset(data, Chr == chr)

\# Fit the loess model with a span based on genomic size

loess_model <- loess(Log2_Ratio ~ Start, data = chr_data, span = span_kb / max(chr_data$Start))

\# Predict smoothed values

chr_data$Smoothed_Log2_Ratio <- predict(loess_model)

\# Append the smoothed results

smoothed_data <- rbind(smoothed_data, chr_data)

}

\# Save the smoothed data

write.csv(smoothed_data, "smoothed_updated_final_result.csv", row.names = FALSE)

\# Example plot for chromosome 1

chr1_data <- subset(smoothed_data, Chr == "chr1")

plot(chr1_data$Start, chr1_data$Log2_Ratio, type = "p", col = "blue",

main = "Chr1 Loess Smoothing", xlab = "Genomic Position", ylab = "Log2_Ratio")

lines(chr1_data$Start, chr1_data$Smoothed_Log2_Ratio, col = "red", lwd = 2)

#------------------------ PCA

############################################

\# This R script performs Principal Component Analysis (PCA)

\# on DNA methylation data for immune cells using genome-wide

\# PMD (Partially Methylated Domains) features.

\# Input:

\#	A tab-delimited text file:

\#	- Rows: Genomic features (e.g., CpGs)

\#	- Columns:

\#	- The first column contains the sample or cell type label ("Cell").

\#	- Remaining columns represent individual samples.

\#	The values are binary:

\#	- 1: the CpG is within a PMD region

\#	- 0: the CpG is outside of a PMD region

\#	- The file should have samples in columns and features in rows;

\#	it will be transposed prior to PCA.

\# Notes:

\#	- The script assumes there are no missing values in the input.

\#	- Manually defined color palette supports up to 18 cell types.

\#	- Axis limits are adjusted dynamically.

############################################

\# Load the data

data_pre <- read.table("input_file.txt", header = TRUE, row.names = 1)

\# Transpose the matrix to get samples in rows and features in columns

data <- t(data_pre)

\# Prepare for PCA: exclude the "Cell" column to retain only numeric values

data_numeric <- data[, -1]

pca_result <- prcomp(data_numeric, scale. = TRUE) # Perform PCA with scaling

\# Convert data to data.frame for downstream processing

data <- as.data.frame(data)

\# Calculate the proportion of variance explained

explained_variance <- (pca_result$sdev^2) / sum(pca_result$sdev^2) * 100

pc1_var <- round(explained_variance[1], 2)

pc2_var <- round(explained_variance[2], 2)

pc3_var <- round(explained_variance[3], 2)

\# Assign colors to each cell type

cell_types <- data$Cell

unique_types <- unique(cell_types)

colors <- rainbow(length(unique_types))

color_map <- setNames(colors, unique_types)

point_colors <- color_map[cell_types]

\# Load ggplot2 for plotting

library(ggplot2)

\# Convert PCA results to data.frame

pca_data <- as.data.frame(pca_result$x[, 1:3]) # Use PC1, PC2, and PC3

pca_data$CellType <- as.factor(cell_types)

\# Define 18 fixed colors manually

color_map <- c("#B3B3B3", "#1B9E77", "#4DAF4A", "#984EA3", "#FF7F00",

"#FFFF33", "#A65628", "#F781BF", "#999999", "#66C2A5",

"#FC8D62", "#8DA0CB", "#E78AC3", "#A6D854", "#FFD92F",

"#E41A1C", "#377EB8")

\# 2D plot: PC1 vs PC2

ggplot(pca_data, aes(x = PC1, y = PC2, color = CellType)) +

geom_point(size = 3) +

scale_color_manual(values = color_map) +

labs(

x = paste("PC1 (", pc1_var, "%)", sep = ""),

y = paste("PC2 (", pc2_var, "%)", sep = "")

) +

theme_minimal() +

theme(

legend.position = "right",

panel.grid.major = element_blank(),

panel.grid.minor = element_blank(),

axis.text = element_text(size = 10),

axis.title = element_text(size = 12),

axis.line = element_line(color = "black", size = 0.1)

) +

coord_cartesian(

xlim = c(min(pca_data$PC1) - 1, max(pca_data$PC1) + 1),

ylim = c(min(pca_data$PC2) - 1, max(pca_data$PC2) + 1)

)

\# 2D plot: PC1 vs PC3 with sample labels

ggplot(pca_data, aes(x = PC1, y = PC3)) +

geom_point(aes(color = CellType), size = 3) +

geom_text(aes(label = rownames(data)), color = "black",

hjust = 0.5, vjust = 0.5, size = 3) +

scale_color_manual(values = color_map) +

labs(

x = paste("PC1 (", pc1_var, "%)", sep = ""),

y = paste("PC3 (", pc3_var, "%)", sep = "")

) +

theme_minimal() +

theme(

legend.position = "right",

panel.grid.major = element_blank(),

panel.grid.minor = element_blank(),

axis.text = element_text(size = 10),

axis.title = element_text(size = 12),

axis.line = element_line(color = "black", size = 0.1)

) +

coord_cartesian(

xlim = c(min(pca_data$PC1) - 1, max(pca_data$PC1) + 1),

ylim = c(min(pca_data$PC3) - 1, max(pca_data$PC3) + 1)

)

#------------------------ heat map and cluster analysis

\# Load the required package

library(pheatmap)

\# Load the data (rows representing TFs, columns representing diseases, with headers and row names included)

data <- read.table(input_file, header = TRUE, row.names = 1, sep = "\t")

\# Convert the data to a numerical matrix

data_matrix <- as.matrix(data)

\# Create the heatmap

pheatmap(data_matrix,

color = colorRampPalette(c("blue", "white", "red"))(100), # Color gradient: blue to white to red

cluster_rows = TRUE,

cluster_cols = TRUE,

show_rownames = TRUE, # Display row names

show_colnames = TRUE, # Display column names

main = "PMD Pattern Heatmap",

breaks = c(seq(0, 1, length.out = 50), seq(1, max(data_matrix), length.out = 51)[-1]), # Breaks at 1 to separate red and blue

legend = TRUE, # Display legend

border_color = NA) # Remove cell borders

#------------------------ Linear regression

############################################

\# This script evaluates the association between hypomethylation index and TCR clonality using linear regression models.

\# Input:

\# - tab-delimited table containing clonality measures and covariates

\### Expected input columns

\# The input table should contain the following variables:
\# TRA_clonality, TRB_clonality, disease_status, sex, age, hypomethylation_index, naive_cd4_index

\### Main analysis steps

\# 1. Load the regression input table

\# 2. Fit linear regression models for TRA and TRB clonality

\# 3. Compare nested models using AIC and adjusted R-squared

\# 4. Extract the regression coefficient, 95% confidence interval, and P value for the hypomethylation index

\# 5. Generate a coefficient plot for the association between hypomethylation index and TRA/TRB clonality

############################################

library(ggplot2)

args <- commandArgs(trailingOnly = TRUE)

if (length(args) != 2) {

stop("Usage: Rscript 04_clonality_regression.R <regression_data.txt> <output_prefix>")

}

input_file <- args[1]

output_prefix <- args[2]

\# Input data

\# Expected columns:

\# TRA_clonality, TRB_clonality, disease_status, sex, age,

\# hypomethylation_index, naive_cd4_index

df <- read.table(

input_file,

header = TRUE,

sep = "\t",

check.names = FALSE,

stringsAsFactors = FALSE

)

\# Model comparison for TRA clonality

fit_tra_1 <- lm(TRA_clonality ~ disease_status + sex + age, data = df)

fit_tra_2 <- lm(TRA_clonality ~ disease_status + sex + age + naive_cd4_index, data = df)

fit_tra_3 <- lm(TRA_clonality ~ disease_status + sex + age + naive_cd4_index + hypomethylation_index, data = df)

\# Model comparison for TRB clonality

fit_trb_1 <- lm(TRB_clonality ~ disease_status + sex + age, data = df)

fit_trb_2 <- lm(TRB_clonality ~ disease_status + sex + age + naive_cd4_index, data = df)

fit_trb_3 <- lm(TRB_clonality ~ disease_status + sex + age + naive_cd4_index + hypomethylation_index, data = df)

\# Save model comparison summary

comparison_table <- data.frame(

model = c("TRA_model1", "TRA_model2", "TRA_model3", "TRB_model1", "TRB_model2", "TRB_model3"),

AIC = c(AIC(fit_tra_1), AIC(fit_tra_2), AIC(fit_tra_3), AIC(fit_trb_1), AIC(fit_trb_2), AIC(fit_trb_3)),

adjusted_R2 = c(

summary(fit_tra_1)$adj.r.squared,

summary(fit_tra_2)$adj.r.squared,

summary(fit_tra_3)$adj.r.squared,

summary(fit_trb_1)$adj.r.squared,

summary(fit_trb_2)$adj.r.squared,

summary(fit_trb_3)$adj.r.squared

)

)

write.table(

comparison_table,

paste0(output_prefix, "_model_comparison.txt"),

sep = "\t",

quote = FALSE,

row.names = FALSE

)

\# Final models including hypomethylation index

fit_tra <- fit_tra_3

fit_trb <- fit_trb_3

\# Extract coefficient, 95% CI, and P value

get_coef_ci_p <- function(fit, term) {

coef_table <- summary(fit)$coefficients

ci <- confint(fit, term)

data.frame(

beta = unname(coef(fit)[term]),

lwr = unname(ci[1]),

upr = unname(ci[2]),

p = unname(coef_table[term, "Pr(>|t|)"])

)

}

term <- "hypomethylation_index"

tra_res <- get_coef_ci_p(fit_tra, term)

trb_res <- get_coef_ci_p(fit_trb, term)

result_table <- rbind(

data.frame(outcome = "TRA_clonality", tra_res),

data.frame(outcome = "TRB_clonality", trb_res)

)

write.table(

result_table,

paste0(output_prefix, "_hypomethylation_association.txt"),

sep = "\t",

quote = FALSE,

row.names = FALSE

)

\# Forest-style coefficient plot

plot_df <- result_table

plot_df$outcome <- factor(plot_df$outcome, levels = c("TRB_clonality", "TRA_clonality"))

p <- ggplot(plot_df, aes(x = beta, y = outcome)) +

geom_vline(xintercept = 0, linetype = 2, color = "grey50") +

geom_errorbarh(aes(xmin = lwr, xmax = upr), height = 0.2, size = 0.6) +

geom_point(size = 2.5) +

theme_classic() +

labs(

x = "Beta (95% CI) for hypomethylation index",

y = NULL,

title = "Association between hypomethylation index and TCR clonality"

)

ggsave(

filename = paste0(output_prefix, "_hypomethylation_association.pdf"),

plot = p,

width = 6,

height = 3.5

)

#------------------------ DMR visualization code

############################################

\# DMR profile plotting

\# This R script visualizes adjusted M values across a genomic region containing a DMR.

\#

\# Input files

\# adjusted_values.tsv

\#	Tab-delimited table containing adjusted M values.

\#	Required columns:

\#	ID : sample identifier

\#	Group1_adj_meanCov : adjusted hypomethylation index

\#	CpG columns : adjusted M values for each CpG probe

\#

\# positions.tsv

\#	Tab-delimited table containing CpG genomic coordinates.

\#	Columns:

\#	CpG chr pos

\#

\# gene annotation

\#	Gene coordinates used for visualization.

\#	Columns:

\#	gene txStart txEnd

\#

\# Output

\#	A plot showing adjusted M values for each CpG site, mean profiles for

\#	top and bottom quartile samples based on hypomethylation index,

\#	the DMR interval, and nearby gene annotations.

############################################

#!/usr/bin/env Rscript

library(data.table)

library(ggplot2)

library(scales)

args <- commandArgs(trailingOnly = TRUE)

if (length(args) != 5) {

stop("Usage: Rscript plot_DMR_profile.R <adjusted_values.tsv> <positions.tsv> <gene_annotation.tsv> <output_prefix> <dmr_chr>")

}

val_file <- args[1]

pos_file <- args[2]

gene_file <- args[3]

output_prefix <- args[4]

dmr_chr <- args[5]

val <- fread(val_file)

pos <- fread(pos_file, header = FALSE)

setnames(pos, c("CpG", "chr", "pos"))

gene_dt <- fread(gene_file)

gene_dt[, mid := (txStart + txEnd) / 2]

required_cols <- c("ID", "hypomethylation_index")

missing_cols <- setdiff(required_cols, names(val))

if (length(missing_cols) > 0) {

stop(paste("Missing required columns in adjusted_values.tsv:", paste(missing_cols, collapse = ", ")))

}

\# Keep CpGs shared between value table and position table

cpgs <- intersect(pos$CpG, setdiff(names(val), required_cols))

if (length(cpgs) == 0) {

stop("No shared CpG columns found between adjusted_values.tsv and positions.tsv")

}

pos2 <- pos[CpG %in% cpgs & chr == dmr_chr][order(pos)]

cpgs_ord <- pos2$CpG

\# Long format: sample x CpG

long <- melt(

val[, c(required_cols, cpgs_ord), with = FALSE],

id.vars = required_cols,

variable.name = "CpG",

value.name = "M_adj"

)

long <- merge(long, pos2, by = "CpG", all.x = TRUE)

\# Define top/bottom quartiles by hypomethylation index

q25 <- quantile(val$hypomethylation_index, 0.25, na.rm = TRUE)

q75 <- quantile(val$hypomethylation_index, 0.75, na.rm = TRUE)

long[, tier := fifelse(

hypomethylation_index <= q25, "bottom25",

fifelse(hypomethylation_index >= q75, "top25", NA_character_)

)]

\# Mean profile for top/bottom quartiles

mean_lines <- long[!is.na(tier),

.(mean_M = mean(M_adj, na.rm = TRUE)),

by = .(tier, chr, pos)

]

mean_lines[, tier := factor(tier, levels = c("bottom25", "top25"))]

\# DMR range from CpG positions shown

dmr_start <- min(pos2$pos, na.rm = TRUE)

dmr_end <- max(pos2$pos, na.rm = TRUE)

\# Plot layout

ymin <- min(long$M_adj, na.rm = TRUE)

ymax <- max(long$M_adj, na.rm = TRUE)

yrng <- ymax - ymin

bar_y <- ymin - 0.08 * yrng

gene_y <- bar_y - 0.10 * yrng

gene_dt[, alt := rep(c("up", "down"), length.out = .N)]

gene_dt[, gene_label_y := ifelse(

alt == "up",

gene_y + 0.08 * yrng,

gene_y - 0.08 * yrng

)]

p <- ggplot(long, aes(x = pos, y = M_adj)) +

geom_point(aes(color = hypomethylation_index), size = 1.2, alpha = 0.75) +

geom_line(

data = mean_lines[tier == "bottom25"],

aes(x = pos, y = mean_M),

linewidth = 0.4,

color = "blue"

) +

geom_line(

data = mean_lines[tier == "top25"],

aes(x = pos, y = mean_M),

linewidth = 0.4,

color = "red"

) +

annotate(

"rect",

xmin = dmr_start, xmax = dmr_end,

ymin = bar_y, ymax = bar_y + 0.03 * yrng,

fill = "darkgreen", alpha = 0.8

) +

annotate(

"text",

x = (dmr_start + dmr_end) / 2,

y = bar_y + 0.06 * yrng,

label = "DMR",

color = "darkgreen",

fontface = "bold"

) +

geom_segment(

data = gene_dt,

aes(x = txStart, xend = txEnd, y = gene_y, yend = gene_y),

inherit.aes = FALSE,

linewidth = 4,

color = "steelblue"

) +

geom_text(

data = gene_dt,

aes(x = mid, y = gene_label_y, label = gene),

inherit.aes = FALSE,

size = 3.2

) +

scale_color_gradient2(

low = "blue",

mid = "white",

high = "red",

midpoint = median(val$hypomethylation_index, na.rm = TRUE),

name = "Hypomethylation index"

) +

scale_x_continuous(labels = comma) +

labs(

x = paste0(unique(pos2$chr), " position"),

y = "Adjusted M value",

title = "DMR methylation profile"

) +

theme_classic() +

theme(

panel.grid = element_blank(),

plot.margin = margin(t = 5.5, r = 5.5, b = 45, l = 5.5)

) +

coord_cartesian(

ylim = c(gene_y - 0.15 * yrng, ymax),

clip = "off"

)

ggsave(

paste0(output_prefix, ".png"),

p,

width = 10,

height = 5,

dpi = 300

)

ggsave(

paste0(output_prefix, ".pdf"),

p,

width = 10,

height = 5

)

***Wiki for custom Python script published on GitHub***

For details, refer to https://github.com/mihshimada/Hypomethylation_immune

#Count Flanking CpG Sites and Sequence Context

CpGContextCounter.py

This script analyzes CpG sites on the DNA methylation array (EPIC, Illumina) and counts the number of additional CpG sites within ±35 bp of the target CpG site. It also extracts the bases immediately flanking the target CpG and classifies them as W (A or T) or S (C or G) based on their identity.

The flanking CpG sites are categorized as follows:

- 0 if there are no additional CpG sites within the flanking region,
- 1 if there is one additional CpG site,
- 2 if there are two additional CpG sites,
- 3 if there are three or more additional CpG sites.

This analysis is performed using the **EPIC_annotation_HeadLines50** sample file (first 50 lines). As long as the first column contains the ID and the second column contains the forward sequence, the analysis can be conducted. The results are output to **EPIC_context.txt**.

\# Calculate the total length of detected DMPs (in Mb)

region_length_summary.py

This script calculates the total genomic length of partially methylated domains (PMDs) using position information output by **methpipe** during PMD prediction.

Each methpipe result file must be in .txt format and contain at least three tab-separated columns: **chr**, **start**, and **end**.

Please replace directory_path with the path to the directory containing the files to be analyzed.

The script processes **all .txt files** in the specified directory as methpipe output files.

Please **do not place unrelated .txt files** in the same directory, as they will also be included in the calculation.

\# Calculate the overlapping regions between PMDs.

pairwise_overlap_length.py

This script compares, in pairs, the overlapping regions between multiple genomic region files (e.g., PMDs predicted by methpipe), calculates their overlapping lengths in megabases (Mb), and outputs the results.

Prepare tab-delimited text files in the target directory that contain at least three columns: **chr**, **start**, and **end**. Please do not place any unrelated text files in that directory.

\# Align and merge multiple bedGraph files into unified genomic intervals

merge_bedgraph.sh

This program merges multiple bedGraph files into a single unified matrix. In our study, bedGraph files were generated from standardized Repli-seq bigWig files—separated by replication timing—using the bigWigToBedGraph tool, a component of the UCSC Genome Browser command-line utilities. These files were then integrated using this program.

project-root/

├── scripts/

│	├── generate_merged_ranges.py	# Step 1: Generate merged genomic ranges

│	└── map_values_to_ranges.py	# Step 2: Map values to merged ranges

├── merge_bedgraph.sh	# Main pipeline script

├── list.txt	# List of input bedGraph files (one per line)

\# Probe-wise Region Annotation Based on Chromosomal Coordinates

annotate_probes.sh

This program annotates each DNA methylation probe on EPIC array with region-specific information (e.g., early/late replication timing) based on chromosomal coordinate overlap.

project-root/

├── CpGRegionAnnotator.py	# Annotate EPIC probes with regional information

├── annotate_probes.sh	# Main pipeline script

├── list.txt	# Text file listing input bedGraph files (one per line)

├── EPIC_context.txt	# Input file containing probe-level information*

├── EPIC_annotation.txt	# Annotation file linking probe IDs to genomic**

*The EPIC_context.txt is intended to be the output file from CpGContextCounter.py, but it is designed to work as long as the ProbeID column is the first column.

**In practice, the program will work as long as the ProbeID is in the first column, the chromosome information is in the third column, and the position information (here based on hg19) is in the fourth column.

\# Merge datasets downloaded from the ENCODE project based on specified criteria.

mergeBEDbyTarget.py

This script processes BED files corresponding to a specific assay target (e.g., CTCF, H3K27ac) in blood samples. It merges overlapping genomic peak regions per chromosome and reports the number of overlapping peaks (i.e., how many input BED files support each merged region).

- Reads a mapping file that links BED file IDs to assay targets.

- Groups BED files by target.

- Merges overlapping peaks across files per target and chromosome.

- Outputs merged regions along with the count of overlapping peaks.

\## Requirements

- BED files in standard 3-column format: `chr`, `start`, `end`.

- A mapping file named `files_assay_target.txt` with two tab-separated columns:

- First column: BED file prefix (e.g., `ENCFF001ABC`)

- Second column: assay target name (e.g., `CTCF`)

project-root/

├── mergeBEDbyTarget.py # Merges overlapping BED regions per target; reports regions that appeared at least once

├── files_assay_target.txt # Tab-delimited file mapping BED file IDs to assay targets (e.g., H3K27ac)

├── A.bed	# Example BED file (input)

├── B.bed	# Example BED file (input)

├── ... # Additional BED files for other assay targets

\# Feature Region Overlap Checker

GenomicFeatureLocator.py

This Python script checks whether genomic features (e.g., CpG sites and SNPs) overlap with annotated regions (e.g., ENCODE regions) across autosomes (chr1–chr22). It outputs a summary of counts and IDs of overlapping features per file.

\## Requirements

- files.txt: A text file listing the ENCODE annotation files to be compared.

Each line should contain one filename (e.g., POLR2A_merged.txt).

- ENCODE Region Files (Assuming the output file from mergeBEDbyTarget.py):

Format: Tab-separated files with the following columns (no header):

Chromosome	start	end

Chromosome must be integers (e.g., 1–22)

Sorted automatically in the script

- feature_file.txt: A feature list file (e.g., CpG sites and SNPs)

Format:

chr1_123456

chr5_234567

...

project-root/

├── GenomicFeatureLocator.py # Identify whether features fall inside/outside regions

├── feature_file.txt	# Mapping between BED file names and assay targets

├── files.txt	# List of merged BED file names to be processed

├── POLR2A_merged.txt	# Example input merged BED file

├── …	# Additional mergedBED files for other assay targets

##Output

- count_result.txt

A tab-separated summary file:

feature	inside	outside

ENCODE1	123	456

ENCODE2	78	921

...

- *_inside_features.txt

For each annotation file, a list of feature IDs that overlapped with any region

\# Summarize CpG information from cis-eQTMs on a per-gene basis

eQTMgeneRanker.py

This program takes the cis-eQTM information reported by Keshawarz et al.* and, for each gene, (i) counts how many CpG sites are linked to that gene through an eQTM relationship, and (ii) assigns each of those CpG sites a rank indicating how strong its association is relative to the other CpGs for the same gene.

*A. Keshawarz et al., Expression quantitative trait methylation analysis elucidates gene regulatory effects of DNA methylation: the Framingham Heart Study. Sci Rep 13, 12952 (2023).

\## Input file format

The script expects a tab-delimited text file (e.g. ALL_cis.tsv) where each row represents one CpG–gene cis-eQTM pair.

Columns are assumed to appear in the following order:

1. rs_ID

CpG probe ID (e.g. Illumina CpG ID such as cg01768446).

2. ProbesetID

Ensembl gene ID (optionally with version), i.e. the gene whose expression is associated with the CpG (e.g. ENSG00000187741.15).

3. RSq

Coefficient of determination (R²) for the eQTM model.

4. Fx

Regression coefficient (effect size) of the CpG on gene expression.

Positive values indicate a positive association; negative values indicate a negative association.

5. T

Test statistic (t-value) for the association.

6. log10P

Base-10 logarithm of the P value for the association (typically negative; more negative = stronger evidence).

7. ENSG_base

Ensembl gene ID without version (e.g. ENSG00000187741).

8. cis_trans

Indicates whether the association is cis or trans (this script uses rows labeled cis).

9. distance_to_TSS

Genomic distance (in bp) from the CpG site to the transcription start site (TSS) of the gene.

10. cg_chr

Chromosome of the CpG site (e.g. 16).

11. cg_pos

Genomic position of the CpG site.

12. gene_chr

Chromosome of the gene/TSS (e.g. chr16).

13. TSS_pos

Genomic position of the transcription start site of the gene.

\# Summarize eQTM information on a per-probe basis

eQTMprobeAggregator.py

eQTMprobeAggregator reorganizes the gene-level eQTM summary into a probe (CpG)-centric view.

It reads the output file produced by the gene-level script (e.g. eQTM_Framingham_cis_1row_gene_rank_NP.txt) and, for each CpG ID, lists all genes with which the CpG forms a cis-eQTM, together with:

the gene label (e.g. ENSG00000123456_P or _N)

the rank of that CpG among all eQTMs for that gene

the number of CpGs in that sign group for the gene

the total number of CpGs linked to that gene

This makes it easy to answer:

“For this CpG, which genes does it regulate, and how strongly is it ranked for each gene?”

\## Input

- File: eQTM_Framingham_cis_1row_gene_rank_NP.txt
- Format: tab-delimited, one line per “gene × sign (P/N)” entry

Required columns (4):

1. GeneID
e.g. ENSG00000123456_P (positive Fx) or ENSG00000123456_N (negative Fx)

2. CpGs(joined)
colon-separated list of CpGs for that gene/sign.
Each CpG is encoded as
cg_id*chr*pos*rank
e.g.
cg00000001*chr1*12345*1:cg00000002*chr1*54321*3

3. #CpGs(sign)
number of CpGs in this sign group (P or N) for that gene

4. #CpGs(total)
total number of CpGs linked to the gene (P + N)
